# No Evidence for a Wakeful Rest Benefit on Associative Memory: A Within-Participant EEG Study

**DOI:** 10.64898/2026.07.01.735795

**Authors:** Marion Walter, Max Lacaze, Samuel Garcia, Nathalie Buonviso, Jane Plailly

## Abstract

Wakeful rest after learning has been proposed to facilitate memory consolidation compared to engaging in a distraction task, with prior EEG studies linking slow oscillation power during rest to better memory performance. However, replication attempts have yielded mixed results. We investigated whether 10 minutes of wakeful rest would enhance associative memory performance relative to 10 minutes of a hippocampus-dependent auditory short-term memory distraction task, using a within-participant design with continuous EEG recording. We employed both a replication-inspired analytical approach, closely modeled on prior work, and a data-specific approach adapted to our dataset. Contrary to our hypotheses, we found no advantage of rest over distraction on associative memory performance. We did, however, observe an order effect: performance was better for the second learning than the first, and this improvement was more pronounced when rest was performed second compared to first. At the neurophysiological level, neither slow oscillation nor alpha power during the post-learning period correlated with memory performance, regardless of analytical pipeline, although cross-over analyses revealed that the choice of EEG reference influenced the direction of some correlations. At the phenomenological level, self-reported mental activity during rest and distraction, as well as trait daydreaming frequency, were not related to memory outcomes, despite the two conditions inducing distinct subjective cognitive states. Together, these findings do not support a robust benefit of post-learning wakeful rest over a hippocampus-dependent distraction task for associative memory, nor do they replicate prior EEG correlates of consolidation. We discuss methodological factors, including task-learning effects in within-participant designs, the coarseness of averaged spectral power measures, and variability in EEG preprocessing pipelines, that may contribute to inconsistencies across the literature, and we call for greater standardization and transparency in future studies.

## 1 Introduction

A growing body of research suggests that short periods of wakeful rest after learning can facilitate memory retention, compared to engaging in distraction tasks. This effect has been observed across diverse memory tasks (e.g., stories (e.g., Dewar et al., 2012; Brokaw et al., 2016; Sacripante et al., 2019; Evans et al., 2021), word lists (e.g., Craig et al., 2014; Dewar et al., 2014; Millar and Balota, 2022; Craig and Greer, 2024), translations (e.g., Mercer, 2015; Wang et al., 2021), picture-word pairs (King and Nicosia, 2022)) and distraction tasks (e.g., spot-the-difference (e.g., Dewar et al., 2012, 2014; Craig et al., 2014, 2016; Sacripante et al., 2019; Martini et al., 2020; Evans et al., 2021), Snood game (Brokaw et al., 2016; Humiston and Wamsley, 2018; Wang et al., 2021). In most studies, resting after learning leads to better recall and less forgetting than being engaged in a distraction task. In a recent systematic review and meta-analysis based on 63 experiments Weng et al. (2025) reported a significant effect of wakeful rest on memory consolidation with a medium effect size. However, these findings are not always consistent accross studies, and a non negligeable amount of them did not find any beneficial effect of rest over distracting task on memory performances (e.g., Fatania and Mercer, 2017; Varma et al., 2017, 2018, 2019; Martini et al., 2018, 2020; Humiston et al., 2019; Tucker et al., 2020; Millar and Balota, 2022). For instance, Brokaw et al. (2016) initially found that 15 minutes of post-learning rest reduced the decline in correct recall and decreased false recall of short stories compared to 15 minutes spent playing the Snood game after learning. However, the same team later failed to replicate these behavioral results (Humiston et al., 2019). Additionnally, in their follow-up meta-analysis of 11 studies investigating the same effect, they found a moderate effect size (Humiston et al., 2019). They concluded that while wakeful rest appears beneficial for memory consolidation, the effect may be smaller than previously believed.

To understand these discrepencies, several lines of research investigated the factors influencing the benefits of wakeful rest on memory consolidation. In an opinion paper, Martini and Sachse (2020) identified the nature of the distraction task as a modulating factor. Indeed, Varma et al. (2017, 2018, 2019) conducted a series of eight experiments using various versions of the N-back task, a working memory paradigm characterized by minimal semantic content and negligeable medial temporal lobe involvement, to systematically manipulate distraction task difficulty, memory content (word-picture pairs and faces) and retrieval type (free recall and recognition). Across all conditions, they found no advantage of wakeful rest over the N-back task on memory performances, suggesting that distraction tasks with limited hippocampal involvement may still be compatible with memory consolidation. Addressing the role of attentional engagement rather than hippocampal involvement, Craig and Greer (2024) showed that post-encoding engagement, and not attentional load (i.e., amount of cognitive resources required by a task), disrupts consolidation: using two difficulties of a Sustained Attention to Response Task (SART) as distractor, they showed that memory scores were higher after rest than after distraction tasks, with no difference between low- and high-attentional load conditions. Dewar et al. (2007) reported equivalent memory impairments across distracting tasks with substantially different content. Their findings showed that interference effects occur regardless of whether the distracting task shares content with the memory task, indicating that content overlap is not a necessary condition for memory interference. Accordingly, when the goal is to examine memory consolidation during wakeful rest rather than content-specific interference, distraction tasks should be designed to minimize overlap with the memory material. Altogether, these findings suggest that for a distraction task to impair memory consolidation through the interruption of active consolidation processes, it should be hippocampus-dependent, capture attention, though not necessarily be difficult, and it should be dissimilar to the learning task. Otherwise, any observed disruption may stem from classical retroactive interference.

Another factor proposed to potentially influence the wakeful rest effect was the amount of mindwandering and daydreaming during wakeful rest. Indeed, Craig et al. (2014) showed that autobiographical thinking cued by sounds, but not listening passively to meaningless sounds, resulted in lower memory performances compared to rest (see also Varma et al., 2018). In the same line, Brokaw et al. (2016) studied the mental activities associated with memory improvements by asking the percentage of time engaged in different mental activities during the post-learning conditions (i.e., autobiographical thinking, thinking about the memory task, meditating, sleeping, thinking about what I am doing). They did not find any link between mental activities during rest and memory performance. However, participants exhibited better memory performances when, during distraction, they spend less time thinking about the distraction task, and more time thinking about their past, current or future life, or meditating. To further study the potential modulatory role of daydreaming and mindwandering on memory consolidation during wakeful rest, Humiston et al. (2019) extended this approach by assessing these tendencies as stable individual traits in everyday life, using validated psychometric instruments: the Daydreaming Frequency Scale (DFS, Stawarczyk et al., 2012), measuring dispositional daydreaming, and the Mindfulness Attention Awareness Scale (MAAS, Brown and Ryan, 2003), measuring trait mindfulness and meditative tendencies. Contrary to their hypothesis, they reported that participants with high propensity of daydreaming and mindwandering showed less memory improvements when learning is followed by rest. This observation opened a new idea in the field: wakeful rest is not sufficient for memory consolidation, and one have to be offline not only from a task to perform, but also from spontaneous thoughts. In this view, wakeful rest could be beneficial for memory consolidation by increasing the probability of experiencing offline states (i.e., strong decrease of externally generated cognition coupled with a strong decrease in internally generated thoughts), or mind blank.

Mechanistically, rest is believed to create a state that supports early memory consolidation, possibly through processes similar to those active during sleep, such as slow oscillations (Wamsley, 2022). To our knowledge, Brokaw et al. (2016) were the first to use electroencephalography (EEG) to explore this idea, showing that slow oscillation (SO, 0.3–1 Hz) power during post-learning rest positively correlated with memory performances. They also showed that alpha (8-12 Hz) power during rest was negatively correlated with memory performances. However, these findings have not been replicated. In a follow-up study, the same team investigated whether brief offline states occurring during a post-learning task could promote memory consolidation. To do so, they operationalized two distinct states: “online” states, characterized by external attentional focus, reduced alpha and SO power, increased pupil diameter, and faster reaction times, were associated with active task engagement; conversely, “offline” states, characterized by increased alpha/SO power, reduced pupil diameter, task-unrelated thoughts, and slower reaction times, were hypothesized to be conducive to memory consolidation. They found that the proportion of time spent in offline states during a post-learning SART significantly predicted subsequent memory performance for a short story. Importantly, this study did not replicate Brokaw et al.’s (2016) findings, in that SO power alone during the SART did not correlate with memory performances, and it did not directly compare rest versus distraction following learning.

In this study, we aimed to replicate the findings from Brokaw et al. (2016) using different memory and distraction tasks, to assess the generalizability of the EEG findings. We compared memory performances to an associative memory task after 10 minutes of wakeful rest or 10 minutes of an auditive short term memory task as distractor with continuous EEG recording in a within-participant design. The choice of the distraction task has been made following the literature recommendation of hippocampal engagement (Borderie et al., 2024). We hypothesized that (1) memory performances after rest would be better than memory performances after distraction, (2) memory performances after rest would be positively correlated with SO power, and negatively correlated with alpha power, during rest. We also investigated the link between mental activities during post-learning, daydreaming frequency, and memory performances.

## 2 Materials and methods

The data analyzed in this article are part of a larger study focusing on the potential modulating impact of the inhalation of a personally pleasant odor on memory consolidation during wakeful rest. The original study includes two groups of participants: a group with an odor diffused during both post-learning conditions, and a group where no odor was presented. Here, we analyzed the data of the no-odor group only. We collected other data not included here. The detailed and complete procedure from the original study is findable in the supplementary file (see original study protocol). This original study has been pre-registered (Walter et al., 2024).

All analyses were implemented using custom Python and R scripts, which are publicly available on GitHub (https://github.com/marion-walter/wakeful-rest-eeg.git) and archived alongside the data on Open Science Framework (https://doi.org/10.17605/OSF.IO/CJDE2, Walter et al., 2026). Comprehensive documentation is included to ensure full reproducibility of the analyses.

### 2.1 Participants

The group consisted of 30 participants in order to replicate the effect of <u>Brokaw et al. (2016)</u> with the same sample size (15 women, 15 men, 29.3 ± 10.5 years, range: 20-57 years). The inclusion criteria were: aged between 18 and 60 years, social security coverage, and willingness to participate in the study. The exclusion criteria were: pregnancy, labor, or breastfeeding, deprivation of liberty by judicial or administrative decision, known cardiovascular, respiratory, olfactory, neurosensory, neurologic or psychiatric disorders, and smoking more than 5 packs a year 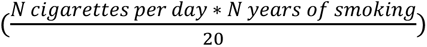. This study was approved on July 2, 2024, by the Ethics Committee (*Comité de Protection des Personnes Ile De France 3*, ID RCB: 2021-A03077-34) in accordance with French regulations for biomedical experiments with healthy volunteers. It was conducted in accordance with the Declaration of Helsinki. Participants were informed clearly and fairly about the study, provided written informed consent, and were compensated for their participation.

### 2.2 Tasks and materials

#### 2.2.1 Stimuli

##### Pictures of items

Fifty pictures of items were used. They were chosen from royalty-free image banks (https://www.pexels.com/fr-fr/ and https://unsplash.com/fr) or generated by artificial intelligence with Microsoft Bing Image Creator (DALL-E 3). They were pictures of animals, musical instruments, plants, food, everyday objects, means of transport, and clothing, displayed on a 1024 × 1024 pixels white background.

Items were selected from an online pilot study to be widely known, easily recognizable, and namable in one unequivocal word. In the pilot 82 pictures were preselected, and 50 volunteers rated the stereotypy of each picture (i.e., how well the image represents what it is supposed to represent, from 0 (not representative at all) to 10 (very representative)) and the familiarity of each picture (i.e., how familiar they were with the item displayed, from 0 (not at all familiar) to 10 (very familiar)), and were asked to name the item with one word, in the most obvious way. We selected the 50 items pictures with the highest scores of stereotypy (8.94 ± 1.55) and familiarity (8.86 ± 1.68), and all of which were named consistently by the volunteers.

##### Pictures of landscapes

Fifty pictures of landscapes were chosen from royalty-free image banks (https://www.pexels.com/fr-fr/ and https://unsplash.com/fr). They were 6000 x 3748 pixels photography of countryside, desert, water, space, forest, underwater, mountains, snowy landscapes, and urban environment.

##### Associations

Fifty associations were built per participant. They consisted of a pair composed of an item picture and a landscape picture. The landscape picture was in the background. The picture of the object was superimposed on that of the landscape, in a central position. Associations were random and differed between participants. For each participant, each picture was used only once. Associations were presented full-screen.

#### 2.2.2 Associative memory task

The memory task itself was adapted from Guinet et al., (2026). It has been coded in Python (version 3.10.19) using Psychopy (version 2023.1.3). The overall structure of the task was as follows: Encoding 1, followed by Initial Retrieval 1, then the first post-learning condition (either Rest or the Distraction task). This was followed by Encoding 2, Initial Retrieval 2, the alternate post-learning condition (i.e., Rest if Distraction was done first, Distraction if Rest was done first), and finally the Surprise Delay Retrieval (Figure 1A). The associative memory task lasted approximately one hour, from Encoding 1 to the Surprise Delay Retrieval.

**Figure 1.**
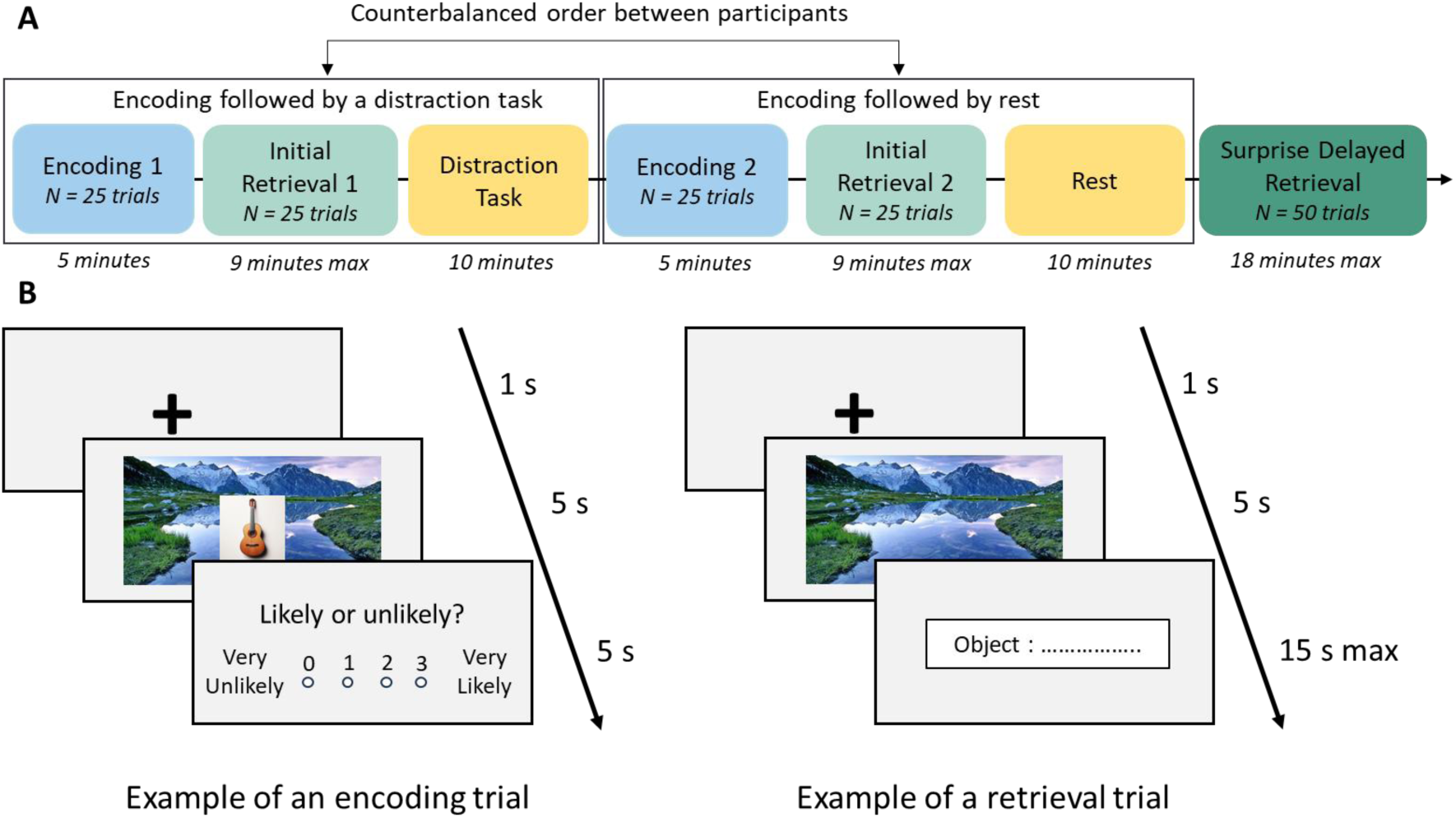
Experimental procedure. A) Associative memory task protocol. Participants completed two learning sessions (Encoding + Initial Retrieval), each followed by either 10 minutes of wakeful rest or 10 minutes of a distraction task, in a counterbalanced order. A Surprise Delayed Retrieval of all the associations concluded the procedure. B) Examples of encoding (left) and retrieval (right) trials. Encoding trial: A 1-second fixation cross was followed by a 5-second display of the landscape-item pair, and a 5-second period to evaluate the likelihood of the association. Retrieval trial: A 1-second fixation cross was followed by a 5-second display of the landscape image, and participants had up to 15 seconds to write the associated item’s name.

##### Encoding

Each Encoding session (Encoding 1, Encoding 2) was composed of 25 trials. A trial consisted in a 1-second presentation of a fixation cross, followed by a 5-second presentation of a landscape-object association. Participants had 5 seconds to rate the likelihood of the association on a discrete scale from 0 (very unlikely) to 3 (very likely) (Figure 1B, left). Participants were instructed to remember as many associations as possible for the retrieval test that followed. An encoding session lasted approximately five minutes.

##### Initial Retrieval

Each Initial Retrieval session (Initial Retrieval 1, Initial Retrieval 2) was composed of 25 trials. A trial consisted of the presentation of one of the landscape pictures presented during the Encoding, for 5 seconds. Then, participants were given up to 15 seconds to type the name of the previously associated item using a keyboard. If they were unable to retrieve the item’s name, they were instructed to enter “0.” A trial ended when the participants pressed the spacebar to validate their answer or when the 15-second time limit elapsed (Figure 1B, right). Order of presentation of the pictures of landscape was random. A retrieval session lasted nine minutes maximum.

##### Surprise Delayed Retrieval

The Surprise Delay Retrieval session was composed of 50 trials similar to the Initial Retrieval sessions, and where the 50 landscapes from Encoding 1 and Encoding 2 were presented in a randomized order. The surprise delayed retrieval lasted 18 minutes maximum.

##### Post-learning rest

The resting session was a period in which participants were instructed to rest quietly, not to fall asleep, keep their eyes open, breathe through their nose, and move as little as possible while sitting on a comfortable chair, for a duration of 10 minutes.

##### Post-learning distraction task

To avoid any interference with the visual material of the associative memory task, as well as to minimize hands and eyes movements for the EEG recording, the distraction task consisted in an auditory task designed by Albouy et al. (2013). It was made of 54 trials. In each trial, participants listened to two six-note melodies (Sequence 1 and Sequence 2) separated by an interval of 1.5 second, and indicated whether the sequences were identical or different by clicking on one of the two responses (Identical or Different). Sequences 1 were generated using MATLAB (R2024a, MathWorks, 2024). under two constraints: each sequence had to contain at least one increasing and one decreasing tone interval, and all sequences had to be unique. For Sequences 2 creation, half of them were modified by altering either the 2^nd^, 3^rd^, 4^th^, or 5^th^ note to a pitch not present in the corresponding Sequence 1. As a result, half of the trials consisted of identical melody pairs, and the other half consisted of mismatched pairs. The task was displayed on a tablet, and lasted 10 minutes. Performances to this task are described and analysed in supplementary file (Supplementary Figure 1).

#### 2.2.3 Questionnaires

##### Daydreaming Frequency Scale

This 12-items scale aims to assess mindwandering frequency in daily life (Stawarczyk et al., 2012). For each item, participants had to choose between 5 possible answers from A (corresponding to the least frequent, scored as a 1) to E (corresponding to the most frequent, scored as a 5). The total score was calculated by summing the scores from each item. A high total score (minimum of 12, maximum of 60) indicated a high frequency of daydreaming.

##### Mental activities questionnaire

We adapted the procedure of Andrews-Hanna et al. (2010) used by (Brokaw et al., 2016), and asked the participants to assess the amount of time engaged in seven predefined categories of mental activities: “Thinking about my past life or imagine my future life”, “Mindwandering”, “Intentionally thinking about the images learned in the memory task”, “Meditating”, “Sleeping”, “Thinking about what I was doing (resting or performing the distraction task)”, “Thinking of nothing, having a mind blank”, “Other: if ticked, specify which mental activity”.

##### Distraction task difficulty questionnaire

Participants had to answer the question “How difficult did you find the task of musical comparison?” (from 0: very easy to 10: very difficult).

The *Daydreaming Frequency Scale* was filled in through LimeSurvey software (LimeSurvey GmbH, n.d.). The *Mental activities questionnaire* and *Distraction task difficulty* question were built using the Python libraries *Dash* (version 3.3.0), and *Psychopy* (version 2025.2.3), respectively.

#### 2.2.4 EEG activity recording

Electric brain activity was recorded with a 64-channel scalp-EEG positioned according to the international 10-20 system using actiCAP nap (actiChamp Plus, Brain Products GmbH, Gilching, Germany) and Brainvision Recorder (Brainvision Recorder, Vers, 1.26.0101, Brain Products GmbH, Gilching, Germany), using Fz as the online reference, and a sampling rate of 500 Hz. The ground electrode was placed in the mid-frontal region. Impedances were kept below 50 kΩ by applying conductive gel to the scalp. Due to technical issues in FCz channel in some participants, this electrode was removed from the analyses for all participants.

### 2.3 Experimental Procedure

The experiment consisted of two sessions a few days apart (median: 3 days, range: 1-28). During the first session (duration: 30 minutes), participants signed an informed consent form, completed the Daydreaming Frequency Scale, and did a picture evaluation procedure not described here (*supplementary file, original study protocol*).

During the second session, participants were equipped with the EEG-cap. A 5-minute baseline recording followed, during which participants were asked to rest, eyes open, breathe through their nose, and move as little as possible. Before starting the associative memory task, instructions were given, and participants underwent a 3-trial training (i.e., an encoding followed by a retrieval) to ensure task understanding. Training trials were the same for all participants. Then, the memory task began (Figure 1A). Participants completed two counterbalanced conditions: a learning (i.e., encoding followed by an immediate retrieval) followed by a 10-minute resting period, and another learning followed by a 10-minute distraction task, resulting in two possible condition orders. After having completed both conditions, participants performed a surprise delayed retrieval of all the associations. EEG was recorded during this whole session.

At the end of the memory task and after being unequipped, participants were asked to complete the mental activities questionnaire, regarding their mental activities during the two post-learning periods: rest and distraction and to complete distraction task difficulty questionnaire. Finally, all the item’s pictures presented to the participants during encoding were displayed one by one, and participants were asked to orally name them as they did during the retrieval phases of the memory task. This step was done to know individual labels of each item and to help scoring. Before the participant left, they were asked if they had any understanding of the objectives and hypotheses of this study. The whole second session lasted approximately 2 hr 30 min.

### 2.4 Data description

#### 2.4.1 Memory performance

For each retrieval trial, participants wrote the name of the item that was associated with the picture of the landscape displayed. Memory performance was scored based on the correspondence between the item presented during encoding and the participant’s response: a score of 1 was assigned for correct retrieval, and 0 for incorrect or omitted responses. Memory performances were then computed by two complementary memory scores.

For each condition (Rest or Distraction), we computed a Global Memory Change score to quantify the change in memory performance between the Initial Retrieval session and the Surprise Delayed Retrieval session. This metric aligns with common practices in the literature, though scoring methods vary across studies (e.g., score at final retrieval – score at initial retrieval, Brokaw et al., 2016; Varma et al., 2017; score at final / score at initial retrieval, <u>Martini et al., 2018</u>, etc.). To enhance precision, particularly in distinguishing between small and large absolute differences (e.g., a change from 3 to 2 vs. 23 to 22), we adopted a refined approach, calculating the score as follows:

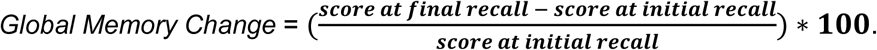

where the score at final retrieval and the score at initial retrieval were the sum of correct responses in the Surprise Delayed and Initial Retrieval, respectively. However, it should be noted that it confuses both trials that were initially retrieved and then forgotten, and trials that were initially forgotten and then retrieved.

Thus, for each condition, we computed a *Relative Memory Change* memory score, which quantifies the change in retention of associations initially learned (as measured during the Initial Retrieval session) by the Surprise Delayed Retrieval session. This score reflects the percentage of initially learned associations that were later retrieved, with a value of 100 indicating a full retrieval. It was calculated as follows:

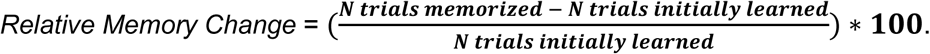

where *N trials initially learned* represents the number of trials correctly retrieved during the Initial Retrieval, while *N trial memorized* refers to the number of trials initially learned that where correctly retrieved during the Surprise Delayed Retrieval.

#### 2.4.2 EEG signals processing

##### Preprocessing

Only EEG data from post-learning conditions (Rest and Distraction) have been analyzed for this article. First, raw EEG data were rescaled to microvolts (µV). To standardize signal duration across participants and conditions, the first 10 seconds of each recording were removed, and the subsequent 10 minutes of data were retained for analysis. To eliminate slow drifts in the signal, a linear detrending process was applied using a high-pass filter with a cutoff frequency of 0.05 Hz, including ringing correction (inspired by De Cheveigné and Arzounian, 2018). Signals were then centered by subtracting the mean value to prepare for Independent Component Analysis (ICA) to exclude ocular and cardiac artifacts. Since ICA performs optimally on signals filtered above 1 Hz, and because our analysis focused on slower oscillations, we avoided removing entire ICA components. This precaution was taken to prevent introducing spectral distortions in frequency bands outside the artifact’s range, which could compromise the integrity of our low-frequency signals of interest. Instead, we employed a targeted approach to remove ocular and cardiac artifacts locally. ICA was first computed on signals filtered between 1 and 100 Hz using the *Infomax* method from the MNE package (version 1.0.2, Gramfort et al., 2013). Ocular and cardiac artifacts were identified using the *iclabel* method for ocular artifacts and the *mne_bad_ecg* method for cardiac artifacts, based on electrocardiogram recordings. These artifacts were then corrected using a combination of Continuous Wavelet Transform and Multistage Singular Spectrum Analysis (CWT-MSSA). The CWT-MSSA algorithm involved: 1) Localizing artifacts using wavelet-based methods; 2) Classifying and correcting blink and cardiac artifacts using Singular Spectrum Analysis or applying a rectification procedure for saccades; 3) Projecting the corrected components back to propagate the correction across all channels. Muscular artifacts were detected using a sliding Root Mean Square threshold window. Artifacted segments were replaced with content matching the spectral properties of the EEG signal (inspired from Ghibaudo et al., 2025). A notch filter was applied to remove line noise at 50 Hz and its harmonics (100, 150, and 200 Hz). Signals were then low-pass filtered at 200 Hz. Remaining large variations in the signals were removed using a threshold based on the Median Absolute Deviation (MAD), following a method inspired by Maddirala and Veluvolu (2021). Finally, signals were mean-centered one last time.

##### Processing

EEG data were processed using two distinct approaches. First, in replication-inspired analyses, to replicate the negative correlation between memory performance and slow oscillation power during rest reported by Brokaw et al. (2016), we adopted their EEG signal processing methodology. Second, in data-specific analyses, we conducted an analysis tailored to our dataset.

For replication-inspired analyses, since Brokaw et al. (2016) recorded EEG signals with mastoid references, we re-referenced our data to the TP9 and TP10 electrodes. For each of the 6 channel, band-averaged power spectral density (PSD) was computed for two frequency bands of interest: slow oscillations (SO, 0.3-1 Hz) and alpha oscillations (8-13 Hz), using Welch’s method with 4-second Hanning windows and a 50% overlap. To account for interindividual variability in EEG amplitude, we used relative power values, normalized such that the total power across all frequency bands summed to a constant value for each analyzed segment. This normalization ensured comparability of spectral power distributions across participants and conditions. Finally, band-averaged PSDs were averaged on participants, conditions and order. One participant was excluded from the Distraction condition due to technical issues during data acquisition.

For the data-specific analyses, we followed the same processing steps described above, with two adjustments. First, the signal from each of the 62 channels was re-referenced to the common average. Second, given that the 4-second windows used in Brokaw et al. (2016) were insufficient to reliably estimate the PSD of slow oscillations — since a 0.3 Hz oscillation has a period of approximately 3.33 seconds — we increased the window length to 20 seconds. This adjustment was based on the estimated period required to capture two adjacent frequency cycles.

### 2.5 Statistical analyses

#### 2.5.1 General information

Descriptive statistics are reported using means and standard deviations (SD) for normally distributed data, and medians and MAD for non-parametric or skewed distributions. Correlations were investigated using Spearman method. For multiple comparisons correction, we used False Discovery Rate (FDR) correction. Results were considered significant at the level of *p* < 0.05. For significant differences effect sizes were reported with Common Language Effect Sizes (CLES) for non-parametric statistical tests (Vargha and Delaney, 2000), and Cohens’ *d* for parametric ones.

Statistical analyses were performed using two approaches: a replication-inspired and a data-specific approaches.

#### 2.5.2 Replication-Inspired Analysis

To align with the methodology of Brokaw et al. (2016), we first performed replication-inspired analyses by computing a memory score inspired from their “change in recall” metric (i.e., *Global Memory Change*). Linear models analyses (i.e., *t*-tests, Analyses of Variance – ANOVAs – or non-parametric equivalents) were conducted using Python language (version 3.11.9), *Pingouin* (version 0.5.5, Vallat, 2018) and *SciPy* packages (version 1.16.3, Virtanen et al., 2020).

##### Memory performances

Linear models were used to assess the main effect of post-learning condition (rest vs. distraction) on both *Global Memory Change* and *Relative Memory Change*. Due to violations of normality in the data and to account for the within-participants design, a non-parametric Wilcoxon test was used. We then tested for a potential order effect of Condition by adding a between-participants Order factor (2 levels: Rest-Distraction, and Distraction-Rest). Because of the mixed-design (within-participant Condition factor and between-participant Order factor) and non-normally distributed data, we used a permutation-based ANOVA to test for an interaction effect between Condition and Order, using the *permuco* package (version 1.1.3, Frossard and Renaud, 2021) with a permutation number set to 5000.

##### Relationship between memory performances and EEG

For each combination of Condition and frequency band, correlations between both *Global Memory Change* and PSD were assessed, using the same parameters as in Brokaw et al. (2016). Therefore, a window duration of 4 seconds was used for PSD estimation, and correlations were computed across the 6 channels used in their study (i.e., F3, F4, C3, C4, O1, O2). For each channel, linear regression was fitted to predict ranked memory performance (*Global Memory Change* and *Relative Memory Change*) with PSD. Correction for multiple comparisons was applied to adjust for the number of channels.

##### Relationship between memory performances and questionnaires

We investigated correlations between memory performances and Daydreaming Frequency Scale scores and mental activities.

#### 2.5.3 Data-specific analysis

In the data-specific analyses, the statistical methods and processing steps were specifically tailored to our experimental design and dataset. Generalized Linear Mixed Models (GLMM) were used, which enabled us to perform trial by trial analysis and to consider participants’ variability as a random effect on non-normally distributed binary data. GLMM were fitted using R language (version 4.5, R Core Team, 2025) on RStudio (Posit team., 2025). Models were fitted using the *glmmTMB* package (version 1.1.13; Brooks et al., 2017), and compared with the *performance* package (version 0.15.2; Lüdecke et al., 2021). Contrasts of interest were examined using the *emmeans* package (version 1.11.2.8; Lenth R and Piaskowski J, 2025). Descriptive model results are reported as estimated marginal means (Standard Error, SE), with 95% confidence intervals (CI95).

##### Memory performances

A GLMM on all trials has been fitted to predict trial-by-trial *performance score* with Condition (Rest vs Distraction) as fixed categorical predictor and a random intercept per participant. This model has been chosen based on model comparison with different random effects structures (see supplementary file for more information about model selection). The outcome variable of the model was ‘memorized’, corresponding to 1 if a trial was memorized between initial and final retrieval, and 0 if a trial was forgotten. Trials that scored 0 on the initial retrieval were not included in this analysis. To test for a potential order effect, we also used a GLMM on all trials with Condition and Order as fixed categorical predictors and a random intercept per participant. We report odds ratios (OR, the ratio of the odds of the outcome occurring in one condition relative the another) with CI95 confidence intervals (CI) and *p*-values to quantify the association between predictors and the outcome.

##### Relationship between memory performances and EEG

Correlations between memory performance and PSD were assessed for each combination of condition and frequency band, using the *Relative Memory Change* metric. A window duration of 20 seconds was used for PSD estimation. Correlations were computed across our full set of 62 channels. For each channel, linear regression was fitted to predict ranked memory performance (*Global Memory Change* and *Relative Memory Change*) with PSD. FDR correction was used to control for the number of channels.

##### Complementary analysis

To check equivalency in memory performance at initial retrieval between (1) conditions (Rest vs Distraction) and (2) learning order (Learning 1 vs Learning 2), we used a permutation-based ANOVA to test for a potential effect of Condition and Learning Order and the interaction between the two on mean score at initial retrieval.

## 3 Results

### 3.1 No beneficial effect of rest on memory performances

Testing for a beneficial effect of the Rest condition on memory performances, we first compared the median *Global Memory Change* (Figure 2A) and *Relative Memory Change* (Supplementary Figure 2) in the Rest and Distraction conditions. Contrary to our hypothesis and the result observed by Brokaw et al. (2016), we observed no beneficial effect of rest on memory performances (*Global Memory Change*: Rest: 0 ± 8.24 % MAD, Distraction: 0 ± 7.61 % MAD; *W* = 134, *p* = 0.46; *Relative Memory Change:* Rest: 5.9 ± 8.75 % MAD, Distraction: 4.26 ± 6.31 % MAD; *W* = 120, *p* = 0.40).

**Figure 2.**
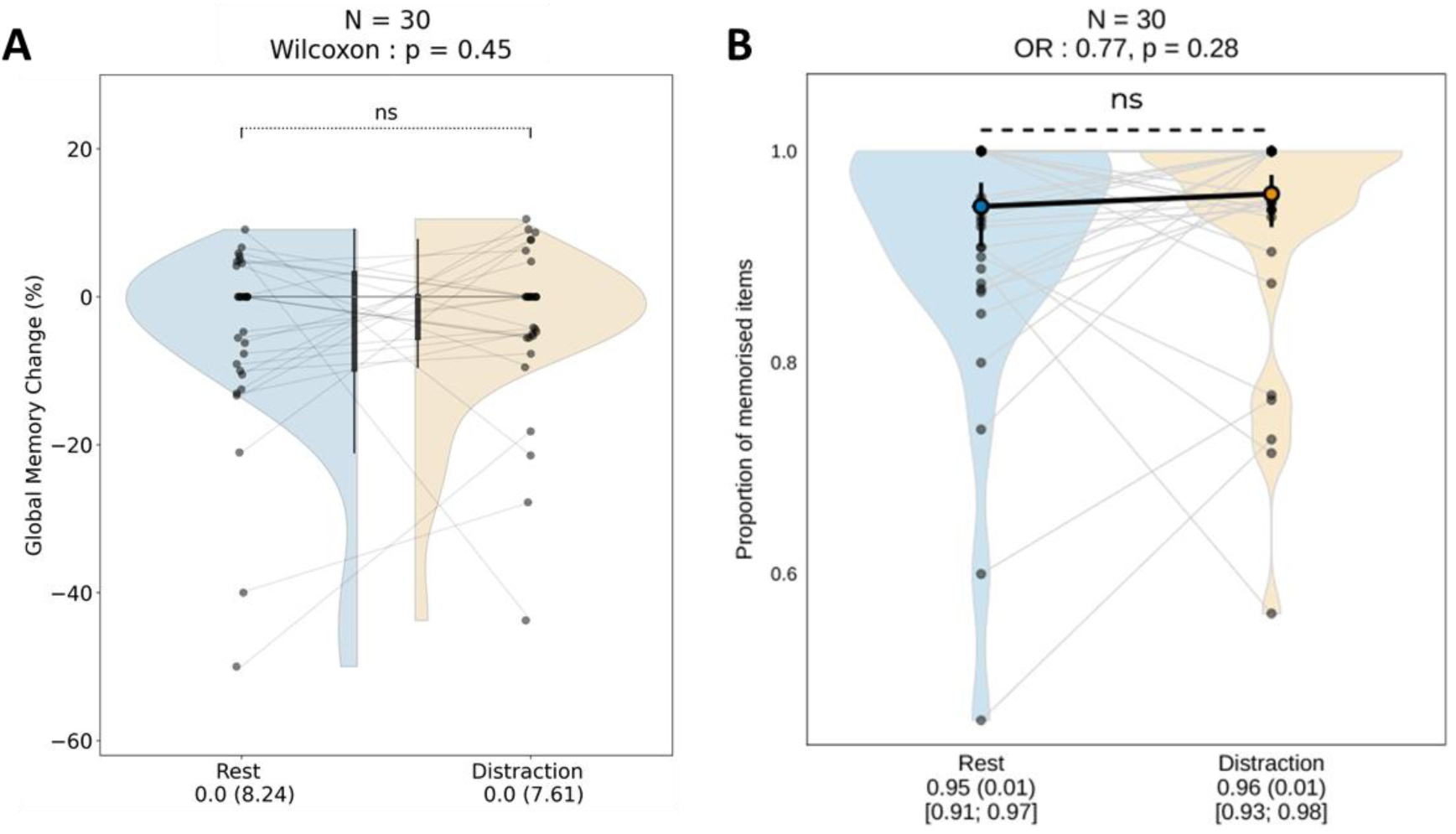
Effects of experimental conditions on memory performances. (A) Distribution of memory scores using *Global Memory Change* for each condition. Inner boxplots indicate central tendencies (whiskers = 1.5 times the inter-quartile range inside beyond the box; box = first quartile, median, third quartile). (B) Distribution of memorized association proportions for each condition. Violin plots show the full data distribution (ranging from minimum to maximum), with individual data points and lines connecting those from the same participant. Estimated marginal means (emmeans) are highlighted with thick dots and lines. ns = non-significant.

Similar results were observed for data-specific analyses (Figure 2B). No significant differences were observed in memory performance between conditions (OR: 0.77, *p* = 0.28). The estimated probability of achieving a score of 1 (i.e., remembering initially retrieved association in the Surprise Delayed Retrieval) in the Rest condition was 0.95 (SE: 0.01, CI95: [0.91, 0.97]), compared to 0.96 (SE: 0.01, CI95: [0.93, 0.98]) in the Distraction condition.

The complementary analysis on initial retrieval only revealed a significant effect of Learning order (*F*(1,28) = 9.13, permutation *p* = 0. 005), with the retrieval performance of the second learning being better than the first, suggesting a learning effect of the memory task. No significant difference was observed between Condition (*F*(1,28) = 0.04, permutation *p* = 0.85) nor in the Learning order x Condition interaction (*F*(1,28) = 0.00, permutation *p* = 1).

### 3.2 No relationship between EEG activity and memory performances

To further investigate the relationship between neural oscillatory activity at rest and memory performances, we examined correlations between SO and alpha power during Rest and subsequent memory outcomes.

The replication-inspired analysis and data-specific analysis both failed to detect a significant correlation between SO power during Rest and memory performance. The Spearman correlation coefficients were *r* = −0.03 (*p* = 0.87) in the replication-inspired analysis (Figure 3A) and *r* = 0.33 (*p* = 0.07) in data-specific analysis (Figure 3B).

**Figure 3.**
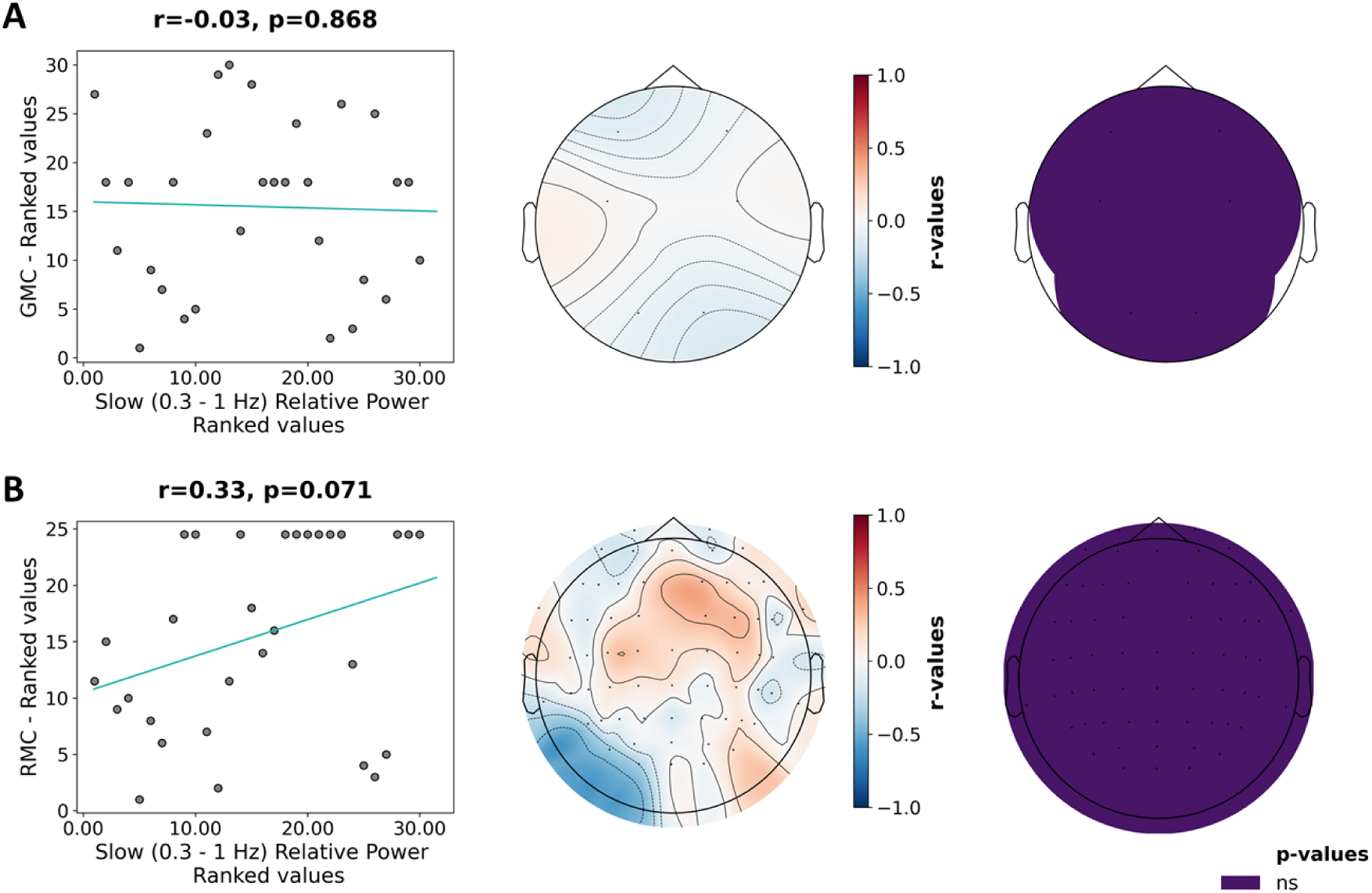
Correlation between slow oscillation power and memory performances for the Rest condition. (A) Replication-inspired analysis restricted to the 6 channels used in Brokaw et al. (2016) with 4-second windows for PSD computation, using TP9 and TP10 channels as references and *Global Memory Change* (GMC) as memory performances (B) Data-specific PSD analysis conducted across 62 channels with 20-second Hanning windows using an average reference and *Relative Memory Change* (RMC) as memory performances. For both analyses: Left panel: linear regression on ranked values of memory performances against slow oscillations (SO, 0.3 – 1 Hz) power. Middle panel: topomap of Spearman correlation coefficients (*r*-values) per channel. Right panel: topomap of channels with significant correlations with memory performances. No significant correlation was observed between SO power and memory performances in either analysis. See supplementary file to see original values of memory performances and SO power (Supplementary Figure 3).

Similarly, no significant correlation was observed between alpha power during Rest and memory performance. The Spearman correlation coefficients were *r* = −0.01 (*p* = 0.95) in the replication-inspired analysis (Figure 4A) and *r* = 0.08 (*p* = 0.66) in data-specific analysis (Figure 4B).

**Figure 4.**
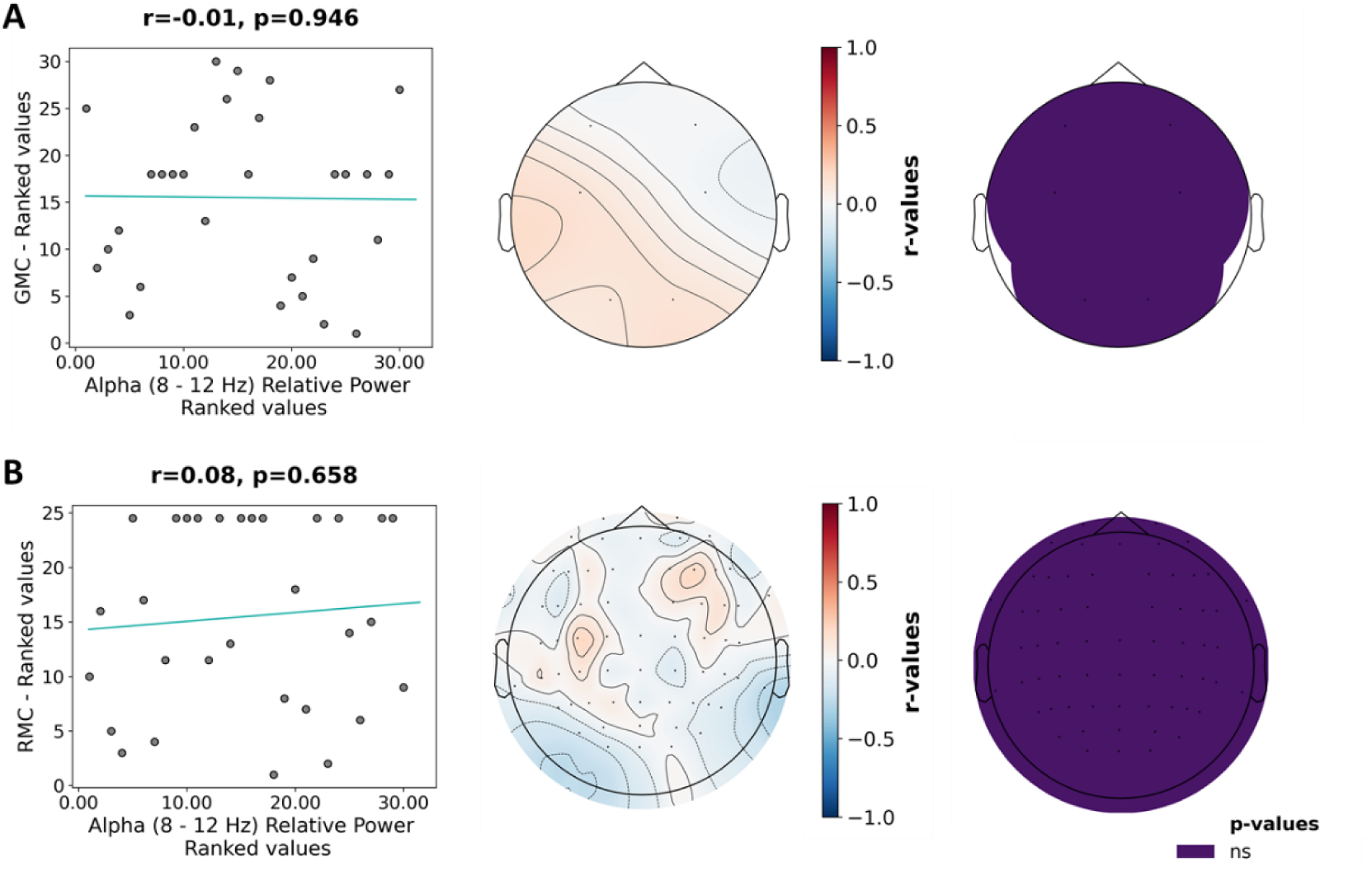
Correlation between alpha power and memory performances for the Rest condition. (A) Replication-inspired analysis restricted to the 6 channels used in Brokaw et al. (2016) with 4-second windows for PSD computation, using TP9 and TP10 channels as references and *Global Memory Change* as memory performances (B) Data-specific PSD analysis conducted across 62 channels with 20-second Hanning windows using an average reference and *Relative Memory Change* as memory performances. For both analyses: Left panel: linear regression on ranked values of memory performances against alpha (8 – 12 Hz) power. Middle panel: topomap of Spearman correlation coefficients (r-values) per channel. Right panel: topomap of channels with significant correlations with memory performances. No significant correlation was observed between alpha power and memory performances in either analysis. See supplementary file to see original values of memory performances and alpha power (Supplementary Figure 4).

Per-channel relative power and its relationship with memory performances across Rest and Distraction are shown in Table 2.

**Table 2.**
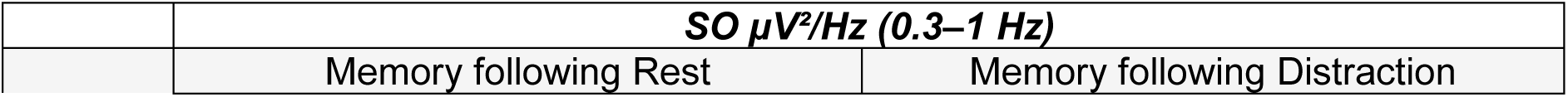
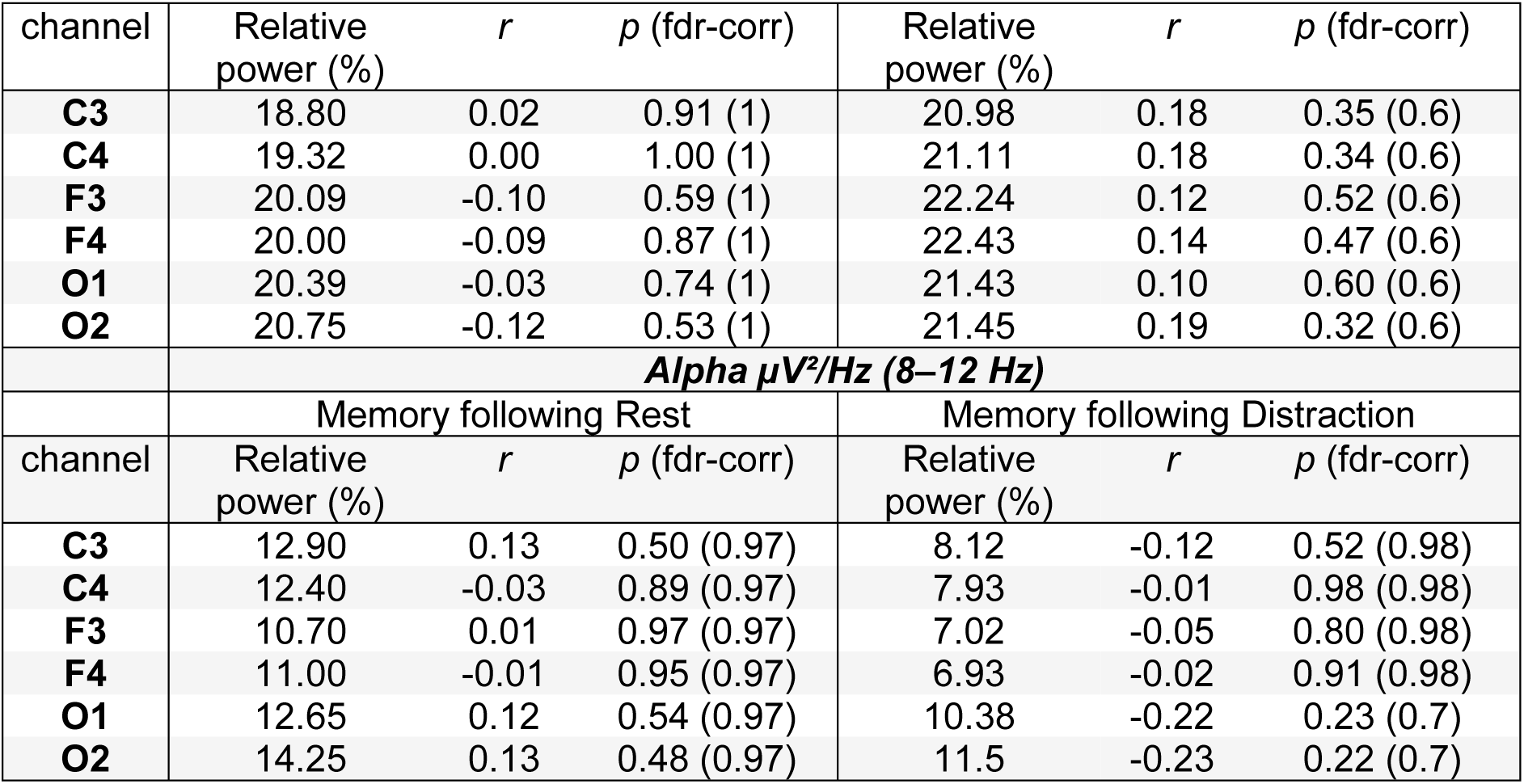
Relationship between EEG activity across Rest and Distraction and memory performances. Spearman correlations assessing the association between EEG power in the Slow Oscillation (SO) and alpha frequency bands, and memory performance (*Global Memory Change*), across Rest and Distraction conditions, and the 6 channels used in the replication-inspired analysis. For each channel, from left to right: relative power in the SO and alpha frequency bands (i.e., sum of normalized PSD within each frequency band, expressed in %), spearman correlation coefficient for each band (*r*), p-values (with adjustment for multiple comparisons using the FDR correction).

### 3.3 Relationship between memory performances, Daydreaming Frequency and mental activities

We examined the relationship between memory performance, Daydreaming Frequency scores, and mental activities during Rest and Distraction. This analysis was motivated by the ambiguous relationship reported in previous studies between these variables. Both *Global Memory Change* and *Relative Memory Change* were used. Because results did not vary between both scores, and because they were highly correlated (*r* = 0.74, *p* < 0.001, Supplementary Figure 5), only *Relative Memory Change* results are described here. Results for *Global Memory Change* are described in supplementary file.

Participants scored in median 43 (± 10.38 MAD, min: 22, max: 59) to the Daydreaming Frequency Scale. There was no significant correlation between *Relative Memory Change* following rest and Daydreaming Frequency scores in the Rest condition (r = −0.14, *p* = 0.47) (Figure 5A).

**Figure 5.**
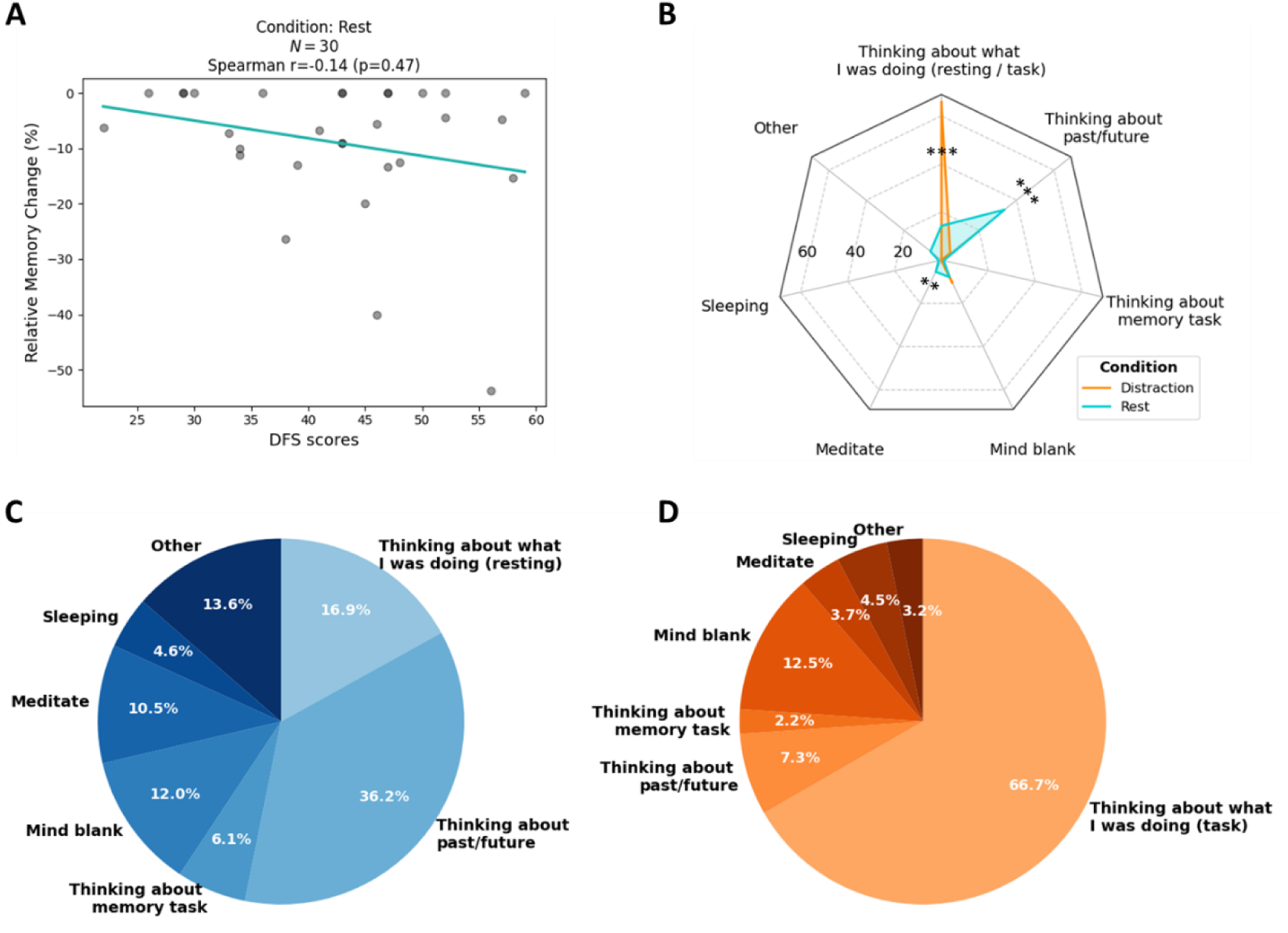
Daydreaming Frequency Scale scores, mental activities, and their relationship with memory performances. (A) Relationship between Daydreaming Frequency Scale (DFS) scores during Rest and memory performance following Rest using *Relative Memory Change*. The blue line represents the regression fit. (B) Comparison of median values of self-reported percentages of the 10-minutes of the post-learning conditions (blue: Rest, orange: Distraction) spent in different mental activities, ** = *p* < 0.01, *** = *p* < 0.001. Proportion of mental activities for the (C) Rest condition and (D) Distraction condition. For visualization purposes, mean values are used instead of median values. Median values and Wilcoxon or paired *t*-test results can be found in Supplementary Figure 6. Note that while Figures 5C and 5D display mean percentages to ensure proportions sum to 100% in the pie charts, the values reported in the text reflect median percentages, consistent with the non-parametric analyses used throughout.

During Rest (Figure 5C), participants reported spending 16.94% of time thinking about what they were doing (i.e., resting) (± 10.67 SD), 1% thinking about associative memory task (± 1.48 MAD), 7.90% having a mind blank (± 11.71 MAD), 5.45% meditating (± 8.08 MAD), 1.15% sleeping (± 1.70 MAD), and 5.80% doing other activities (± 8.60 MAD). The “other activities” category most frequently included: planning daily/weekend activities, observing the surroundings, thinking about the experiment, hunger-related thoughts, mental repetition of songs, time monitoring, awareness of nasal breathing, imaginative scenarios, and stress-related thoughts. Notably, when participants reported “sleeping” (N = 14) all clarified that they were experiencing drowsiness or resisting sleep rather than actually falling asleep. During Distraction (Figure 5D), participants reported spending in average 66.67% of time thinking about what they were doing (i.e., being focused of the task) (± 19.54 SD), 4.70% thinking about past/future (± 6.97 MAD), 0% thinking about associative memory task (± 0 MAD), 10.55% having a mind blank (± 7.93 MAD), 0% meditating (± 0 MAD), 0% sleeping (± 0 MAD), and 0% doing other activities (± 0 MAD). The “other activities” category most frequently included: thinking about the experiment, mental repetition of songs, mindwandering and unrecallable thoughts.

Participants reported spending significantly more time thinking about past/future in Rest compared to Distraction (Figure 5B; *W* = 3.50, *p* < 0.001, CLES = 0.89). A similar pattern was observed for meditating (W = 34.00, *p* < 0.01, CLES = 0.67). Conversely, participants spent significantly more time focusing on what they were doing in the Distraction condition compared to Rest (*t* = 11.85, *p* < 0.001, Cohen’s *d* = 3.19). No significant differences were observed for thinking about memory task (i.e., rehearsal), sleeping, nor having a mind blank (*W*’s > 65; *p*’s > 0.08). No correlation was found between mental activities during Rest and Distraction and *Relative Memory Change* (*r*’s < |0.30|; corrected *p*’s > 0.24; Supplementary Figure 7).

### 3.4 Order effect

Given that previous studies have reported a significant effect of condition order on memory performances (Brokaw et al., 2016; Martini et al., 2018), we investigated the interaction between Condition (Distraction, Rest) and Order (First, Second) using both the replication-inspired analyses and data-specific analyses.

For the replication-inspired analyses, the evolution of memory scores between initial and surprise delayed retrieval (*Global Memory Change*) did not significantly differ between Condition (*F*(1,28) = 0.35, permutation *p* = 0.57) or Order (*F*(1,28) = 1.45, permutation *p* = 0.25) nor Condition and Order interaction (*F*(1,28) = 3.27, permutation *p* = 0.08).

For data-specific analyses, contrasts analyses revealed a significant interaction between Condition and Order (Figure 6). Post-hoc comparisons revealed that the remembering of the learned associations was always higher for the second learning than for the first (Order Distraction-Rest, Rest: 0.98, SE: 0.01, CI95: [0.95, 0.99], Distraction: 0.95, SE: 0.02, CI95: [0.89, 0.98]; OR: 2.71, *p* = 0.02; Order Rest-Distraction, Rest: 0.91, SE: 0.03, CI95: [0.82, 0.96], Distraction: 0.97, SE: 0.01, CI95: [0.93, 0.99]; OR: 0.30, *p* < 0.001). Moreover, when comparing each Condition between Orders, post-hoc tests revealed that memory performances for Rest were significantly higher when Rest was performed second compared to when it was performed first (OR: 4.8, *p* = 0.01). Memory performances for Distraction did not differ significantly between Order (OR: 0.53, *p* = 0.31). Finally, when comparing conditions for each learning, no comparisons were significant (first learning, OR: 1.78, *p* = 0.31; second learning, OR: 1.45, *p* = 0.57). Our results showed a clear advantage of the second learning, and even more when considering Rest than Distraction condition.

**Figure 6.**
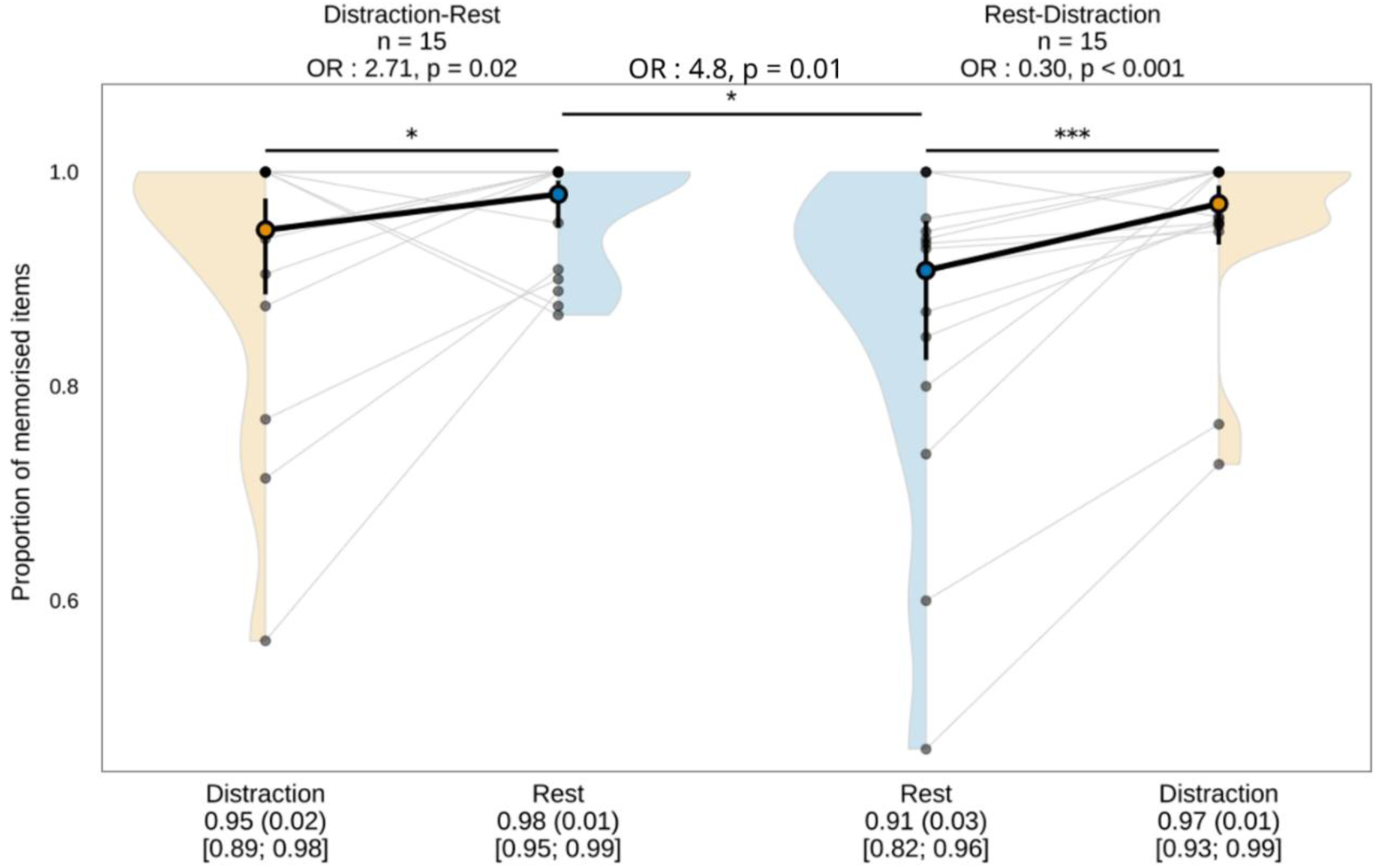
Effects of order of experimental conditions on memory performances. Distribution of percentage of memorized items for each condition (Distraction in yellow, Rest in blue), within each condition order. Left panel: Distraction – Rest, Right panel: Rest – Distraction. Data points and violin plots indicate real data, with lines connecting those from the same participant, while estimated marginal means (emmeans) are highlighted with thick dots and lines. * = *p* < 0.05, *** = *p* < 0.001.

## 4 Discussion

Our goal was to test the beneficial effect of short periods of wakeful rest after learning for memory retention, compared to engaging in distraction tasks, and the potential role of slow brain oscillations in this consolidation process during rest. We used a within-participant design to assess memory performance in an associative memory task followed either by 10 minutes of wakeful rest or by a 10-minute distraction task, both accompanied by continuous EEG recording. Our analytical approach combined a replication-inspired method, closely aligned with Brokaw et al. (2016), and a data-specific method, tailored to our specific dataset. Contrary to our hypothesis, we found no beneficial effect of wakeful rest on memory performances, compared to Distraction condition. However, we found an advantage of the second learning compared to the first learning, and even more when considering Rest than Distraction condition. On the neurophysiological level, we found no relationship between EEG slow oscillations or alpha power, and memory performances. On the phenomenological level, we also found no significant relationship between mental activity during Rest and Distraction, daydreaming frequency, and memory performances.

### 4.1 No replication of the wakeful rest effect on memory performances

The present study found no beneficial effect of post-learning wakeful rest on memory performance compared to a distraction task. We propose a set of methodological and theoretical explanations that, taken together, highlight the complexity of studying consolidation during wakefulness.

While rest had been demonstrated to be beneficial for memory consolidation in several studies, (e.g., Gottselig et al., 2004; Dewar et al., 2012, 2014; Craig et al., 2014, 2016; Mercer, 2015; Brokaw et al., 2016; Schlichting and Bäuml, 2017; Sacripante et al., 2019; Evans et al., 2021; Wang et al., 2021; King and Nicosia, 2022; Craig and Greer, 2024), and reported in a recent systematic review and meta-analysis based on 63 experiments with a medium effect size (Hedges’s *g* = 0.448, 95% CI [0.339, 0.557], *p* < .001; Weng et al., 2025) our result is consistent to a subset of studies that also failed to replicate the wakeful rest benefit. Varma et al., (2017) attributed their negative result to the nature of their distraction task (i.e., N-back task) arguing it did not engage the hippocampal, thought to interfere with consolidation. Although our distraction task is hippocampo-dependant (Borderie et al., 2024), it may rely on different hippocampal subsystems than those engaged during the associative encoding phase, potentially limiting its interference. Humiston et al. (2019), who failed to replicate the behavioral findings from Brokaw et al. (2016) using an identical protocol, invoked an insuficient sample size or sampling variability. Finally, studies attempting to extend the wakeful rest paradigm to ecological settings have consistenly found no benefit (Brandmark et al., 2020; Leetham et al., 2024; Seban et al., 2026), underscoring the difficulty of translating laboratory findings to real-world situations.

A central issue in wakeful rest research is that the condition itself is poorly standardized. Participants left to rest wakefully do not share a common cognitive or mental state: some engage in passive visual exploration, others ruminate on personal concerns, and others struggle against drowsiness. To a lesser extent during the distraction task, even though the state of the participants was constrained by the task, they still experienced fluctuations in their attentional engagement and free thoughts that were not monitored online. These variable inner experiences could substantially modulate consolidation processes, contributing to the large inter-individual variability observed in memory performance across studies. This suggests that the post-learning condition may be a poor proxy for the underlying neural processes relevant to awake consolidation.

In the present study, we attempted to characterize mental activity during both conditions via post-experiment questionnaire, like it has been generally done in previous studies. Participants reported spending significantly more time engaged in autobiographical thoughts during the Rest condition, and more time thinking about the task they were doing during the Distraction condition confirming that the two post-learning conditions induced significantly distinct cognitive states. However, we found no significant correlation between self-reported mental activity and memory performance. This absence of a relationship aligns with the mixed results in the existing literature: Brokaw et al. (2016) reported a positive correlation between mindwandering and memory performance during Rest, Humiston et al. (2019) found a negative correlation, Wamsley and Summer (2020) observed associations that depended on the type of memory task, and Wang et al. (2021) found no relationship. The inconsistency across studies suggests that the link between subjective mental states and consolidation is neither direct nor robust, and may depend on unmeasured moderating variables. More broadly, and consistent with concerns raised in prior studies, retrospective estimation of the time spent in different mental states across a 10-minute window is inherently difficult and subject to systematic distortion, limiting the precision and reliability of these self-reports. Future studies may benefit from using experience sampling or continuous ratings to capture mental activity in a more temporally resolved manner. In this regard, more precise analyses incorporating physiological and behavioral indices of attentional engagement, such as pupillometry or response time variability alongside subjective ratings of attentional focus (Wamsley and Summer, 2020; Wamsley et al., 2023), may be necessary to uncover the subtle dynamics underlying consolidation during wakefulness.

Finally, we observed a strong memory performance difference between participants, emphasizing the importance of a within-participant design, despite the risk of a task-learning effect. Indeed, the scores for *Global Memory Change* ranged from minimum −50 to maximum 9.09 in the Rest condition and from minimum −43.75 to maximum 10.53 in the Distraction condition. This strong disparity can of course be due to normal inter-participant variability. However, based on participants oral accounts during the final interview, the encoding strategy used by participants seems to play a decisive role in memory performances. Indeed, participants who reported creating short stories to build associations between landscape-object picture pairs generally showed better memory performance. Although our study did not focus on encoding strategies, this factor could potentially influence the effect of rest during wakefulness and warrants further investigation in future research, thereby highlighting the difficulty of designing a memory task that allows for equivalent encoding among participants, regardless of strategy.

The literature as a whole underscores the need to more systematically incorporate null results, to identify potential moderating variables, and to better account for methodological variation across studies (Martini and Sachse, 2020b; Weng et al., 2025). Parameters such as participants age, with older adults benefiting more than younger participants (Martini et al., 2019), memory tests, with recall measures being more sensitive than recognition (Millar and Balota, 2022; Weng et al., 2025), and memory tasks (Craig et al., 2016) have all been identified as modulators of the wakeful rest effect,, and. Taken together, these factors highlight the complexity of isolating the wakeful rest benefit and the importance of rigorous, transparent reporting of methodological choices.

### 4.2 Absence of EEG correlates of memory consolidation

We found no significant correlation between EEG activity during post-learning conditions and subsequent memory performances. Surprisingly, in our data, relative SO power was higher during the Distraction condition than during Rest (Table 2). While this difference was modest, it raises questions about the specificity of SO power as a predictor of memory consolidation and suggests that absolute power levels may not straightforwardly reflect the quality of consolidation processes. This discrepancy aligns with mixed findings in the literature: while some studies (e.g., Brokaw et al., 2016; Bencze et al., 2025) report a link between SO power during rest and long-term memory, others, like Wang et al. (2021), found no such correlation across multiple frequency bands, despite better memory outcomes for rest and nap conditions compared to distraction. Several factors may explain these inconsistencies, such as protocol differences (i.e., memory tasks), retention delays or measurement approach (immediate recall test and average of spectral power in Brokaw et al., 2016, Wang et al., 2021 and our study; recall test after 7 days, and change in spectral power from a pre-learning baseline to the post-learning rest period in Bencze et al., 2025). However, we also note a more fundamental issue: averaging spectral power across an entire 10-minute rest epoch and correlating it with a single memory score may be a too coarse approach to capture the subtle and transient neurophysiological dynamics that support consolidation.

According to the two-stage model of memory (Buzsáki, 1989), new experiences are initially encoded in parallel across both the hippocampus and neocortical networks. The hippocampus subsequently reactivates these memory traces during offline states, most notably during sleep, but potentially also during wakeful rest (Mednick et al., 2011; Wamsley, 2022), orchestrating their replay in the neocortex. This iterative reactivation is thought to underlie the stabilization of initially fragile memory traces into durable long-term representations. Critically, the hippocampal-neocortical dialogue is believed to rely on precise coupling between fast and slow oscillatory rhythms (Schreiner and Staudigl, 2020), SO acting as the primary pacemaker. During SO, cortical networks alternate between hyperpolarized down and depolarized up states, each lasting from 0.2 to 1 second. Hippocampal ripples and thalamic spindles consistently align with SO up states during slow-wave sleep (SWS), giving rise to a fine-tuned cross-frequency coupling widely recognized as an important electrophysiological substrate of memory consolidation (Clemens et al., 2007; Staresina et al., 2015; Keeble et al., 2025).

While this mechanism is well-established during SWS, the functional role of SO during wakefulness remains considerably less clear. Although SOs have been observed in humans during wakefulness (Demanuele et al., 2010), the oscillatory landscape of the waking brain differs markedly from that of SWS, suggesting that the consolidation mechanisms at play may be partially distinct (Wamsley, 2022). Scalp-recorded SO power during brief rest periods may provide only an indirect and potentially incomplete index of the consolidation processes of interest. Recent work using intracranial EEG in pharmaco-resistant epileptic patients has provided direct access to recordings of deep brain structures such as the hippocampus and medial temporal lobe, enabling direct observation of ripple activity and memory replay during wakefulness (Axmacher et al., 2008; Zhang et al., 2018; Sjøgård et al., 2025; Causse et al., 2026). Complementing these findings, magnetoencephalography studies in healthy participants have identified replay-like neural activity during brief periods of wakeful rest, and the extent of this replay predicts subsequent skill consolidation (Buch et al., 2021). Together, these approaches offer a promising avenue for bridging the gap between mechanistic evidence from animal models and the correlational findings typically obtained with scalp EEG in humans.

### 4.3 The need for methodological transparency

Our analytical approach combined a replication-inspired method, closely aligned with Brokaw et al. (2016), and a data-specific method, tailored to our specific dataset. At the behavioral level, results were highly consistent across analyses, regardless of whether Global Memory Change or Relative Memory Change was used as the memory score, and whether linear models (replication-inspired approach) or trial-by-trial GLMMs (data-specific approach) were employed. This convergence across methods strengthens confidence in the robustness of our behavioral findings.

At the neurophysiological level, however, notable differences emerged between the two analytical pipelines. Neither the replication-inspired analysis (Global Memory Change, mastoid-like reference, 6 channels, 4-second Hanning windows for PSD computation) nor the data-specific analysis (Relative Memory Change, average reference, 62 channels, 20-second Hanning windows) yielded significant correlations between SO power and memory performance. Yet, differences in effect sizes and *p*-values were observed between pipelines. To disentangle the respective contributions of the memory score and the EEG preprocessing parameters in these discrepancies, we conducted exploratory cross-over analyses to explore the impact of reverting the two memory scores, as well as window duration for PSD computation, and the offline reference on the correlation between memory performances and SO power. In short, while reverting the memory scores and the window duration did not seem to change the correlations values, the offline reference change almost reverts the direction of some correlations though it did not reach the threshold of statistical significance. Further details on these analyses are available in the supplementary file (Supplementary Figure 8, 9 and 10).

Taken together, these observations highlight a broader issue in the field: the considerable variability in neurophysiological analysis pipelines across studies, often accompanied by insufficient reporting and justification of parameter choices.

### 4.4 Order effects interpretation

To our knowledge, only a minority of within-participant studies have explicitly tested for order effects. Evans et al. (2021) found no significant effect of Order nor interaction between Condition and Order on memory performances. Brokaw et al. (2016) and Martini et al. (2018), just as our results showed, both reported better memory performance when Rest was completed second rather than first. Notably, in both studies, memory performance in the Distraction condition did not improve when that condition was presented second. Brokaw et al. (2016) attributed this pattern to a facilitatory effect of distraction on subsequent new encoding, while Martini et al. (2018) argued that distraction interferes with the consolidation of material learned prior to the rest period. We propose an alternative interpretation: the observed order effect may primarily reflect a task-learning effect, whereby participants became more familiar with the associative memory paradigm and developed more effective encoding strategies during the second attempt. The apparent advantage of Rest when presented second may not stem from condition-specific consolidation dynamics, but rather from the interplay between task familiarity (associative memory task is easier when performed the second time than the first) and emotional state (e.g., confidence of anxiety-related thoughts, may be less present during Distraction due to task focus).

A critical interpretive challenge in within-participant designs is the difficulty of dissociating a post-encoding consolidation effect from a learning-related improvement already present at the time of encoding. To address this, we examined initial retrieval performance alone, prior to any post-learning condition, and found that the performance advantage for the second learning was already evident at this stage. This result constitutes an additional argument in favor of a task-order learning effect, independent of any consolidation process. To our knowledge, such a learning effect has not been explicitly reported in wakeful rest memory literature using within-participant designs, though its absence from published reports may reflect a lack of systematic investigation rather than an absence of the phenomenon. This raises a broader methodological concern: within-participant designs, while statistically efficient and well-suited to control for individual differences, may be particularly vulnerable to order effects when the target effect is of small to moderate magnitude. In such cases, task learning may obscure, dilute, or interact with the consolidation effect of interest, complicating the interpretation of condition differences.

### 4.5 Conclusion

The present study failed to find a significant advantage of post-learning Rest over Distraction on associative memory performance, nor any relationship between post-learning SO or alpha power, mental activities, and subsequent memory performances. Although our aim was to conceptually replicate and assess the generalizability of the behavioral and EEG findings of Brokaw et al. (2016), several methodological differences may account for the discrepancy, including the use of an associative memory task with cued retrieval rather than story recall with free recall, a hippocampus-dependent auditory distractor task, and a delayed final retrieval test administered after both conditions to preserve the surprise effect and limit intentional rehearsal. Despite these differences, our protocol followed widely used procedures in the wakeful rest literature and was grounded in prior evidence. Our cross-analysis comparisons further highlight that small-to-moderate effects are particularly sensitive to EEG preprocessing decisions, notably the reference choice and PSD estimation window, underscoring the importance of methodological transparency in this literature. Whether the absence of a wakeful rest benefit reflects differences between paradigms, insufficient sensitivity of broadband SO power as a consolidation index, or uncontrolled inter-individual factors such as encoding strategy, remains an open question. Future work should prioritize more precise neurophysiological measures, systematic reporting of order effects, and greater standardization of analytical pipelines to build a more cumulative science of offline memory consolidation.

## 5 Acknowledgements

We would like to thank D^re^ Anne Caclin for providing us with the short-term auditory memory task that we used as the distraction task (including all the necessary materials for administering it) and for her help in adapting it to our needs; Marie Touzet—Robin and Lauryne Zacharia for their help in designing the study and collecting data; Belkacem Messaoudi for the technical support during the experimentation; Christelle Daudé for her help with participant recruitment and EEG setup; Iris Guggenbuhl for her help with the EEG setup; Guillemette Colas des Francs for her participation in mental activity analysis; D^r^ Valentin Ghibaudo for his help and advices on Python scripts and analyses; Maël Delem for his help with generalized linear mixed-effects model analysis and R scripts.

## 8 Authors contributions

**Marion Walter:** Conceptualisation, Methodology, Investigation, Data curation, Formal analysis, Software, Visualisation, Writing – Original draft Preparation. **Max Lacaze:** Data curation, Software, Formal analysis**. Samuel Garcia:** Software**. Nathalie Buonviso:** Conceptualisation, Supervision, Funding acquisition, Writing - Review & Editing. **Jane Plailly:** Conceptualisation, Supervision, Writing - Review & Editing.

## 9 Data, scripts, code, and supplementary information availability

Data are available online: https://doi.org/10.17605/OSF.IO/CJDE2; Walter et al. (2026) Scripts and code are available online: https://doi.org/10.17605/OSF.IO/CJDE2; Walter et al. (2026)

Supplementary information is available online: https://doi.org/10.17605/OSF.IO/CJDE2; Walter et al. (2026)

## References

actiChamp Plus [Apparatus] (2026) Gilching, Germany: Brain Products GmbH.

Albouy P, Mattout J, Bouet R, Maby E, Sanchez G, Aguera P-E, Daligault S, Delpuech C, Bertrand O, Caclin A, Tillmann B (2013) Impaired pitch perception and memory in congenital amusia: the deficit starts in the auditory cortex. Brain 136, 1639–1661. 10.1093/brain/awt082

Andrews-Hanna JR, Reidler JS, Huang C, Buckner RL (2010) Evidence for the Default Network’s Role in Spontaneous Cognition. J Neurophysiol 104, 322–335. 10.1152/jn.00830.2009

Axmacher N, Elger CE, Fell J (2008) Ripples in the medial temporal lobe are relevant for human memory consolidation. Brain 131, 1806–1817. 10.1093/brain/awn103

Bencze D, Marián M, Szőllősi Á, Simor P, Racsmány M (2025) Increase in slow frequency and decrease in alpha and beta power during post-learning rest predict long-term memory success. Cortex 183, 167–182. 10.1016/j.cortex.2024.11.012

Borderie A, Caclin A, Lachaux J-P, Perrone-Bertollotti M, Hoyer RS, Kahane P, Catenoix H, Tillmann B, Albouy P (2024) Cross-frequency coupling in cortico-hippocampal networks supports the maintenance of sequential auditory information in short-term memory. PLOS Biol 22, e3002512. 10.1371/journal.pbio.3002512

BrainVision Recorder (Vers. 1.26.0101) [Software] (2026) Gilching, Germany: Brain Products GmbH.

Brandmark A, Byrne M, O’Brien K, Hogan K, Daniel DB, Jakobsen KV (2020) Translating for Practice: A Case Study of Recommendations From the Wakeful Rest Literature. Teach Psychol 47, 92–96. 10.1177/0098628319889268

Brokaw K, Tishler W, Manceor S, Hamilton K, Gaulden A, Parr E, Wamsley EJ (2016) Resting state EEG correlates of memory consolidation. Neurobiol Learn Mem 130, 17–25. 10.1016/j.nlm.2016.01.008

Brooks ME, Kristensen K, Benthem KJ van, Magnusson A, Berg CW, Nielsen A, Skaug HJ, Mächler M, Bolker BM (2017) glmmTMB Balances Speed and Flexibility Among Packages for Zero-inflated Generalized Linear Mixed Modeling. The R Journal 9, 378–400. https://digitalcommons.unl.edu/r-journal/675/

Brown KW, Ryan RM (2003) The benefits of being present: Mindfulness and its role in psychological well-being. J Pers Soc Psychol 84, 822–848. 10.1037/0022-3514.84.4.822

Buch ER, Claudino L, Quentin R, Bönstrup M, Cohen LG (2021) Consolidation of human skill linked to waking hippocampo-neocortical replay. Cell Rep 35, 109193. 10.1016/j.celrep.2021.109193

Buzsáki G (1989) Two-stage model of memory trace formation: A role for “noisy” brain states. Neuroscience 31, 551–570. 10.1016/0306-4522(89)90423-5

Causse AA, Curot J, Lopes-dos-Santos V, Nunes-da-Silva R, Barron HC, Dornier V, Denuelle M, De Barros A, Sol J-C, Lotterie J-A, Lehongre K, Fernandez-Vidal S, Frazzini V, Navarro V, Valton L, Barbeau EJ, Denison T, Reddy L, Dupret D (2026) A learning-evoked slow-oscillatory architecture paces population activity for offline reactivation across the human medial temporal lobe. Neuron, S0896627326003752. 10.1016/j.neuron.2026.05.004

Clemens Z, Mölle M, Eross L, Barsi P, Halász P, Born J (2007) Temporal coupling of parahippocampal ripples, sleep spindles and slow oscillations in humans. Brain 130, 2868–2878. 10.1093/brain/awm146

Craig M, Greer J (2024) Post-encoding task engagement not attentional load is detrimental to awake consolidation. Sci Rep 14, 12345. 10.1038/s41598-024-53393-6

Craig M, Sala SD, Dewar M (2014) Autobiographical Thinking Interferes with Episodic Memory Consolidation. PLOS ONE 9, e93915. 10.1371/journal.pone.0093915

Craig M, Wolbers T, Harris MA, Hauff P, Della Sala S, Dewar M (2016) Comparable rest-related promotion of spatial memory consolidation in younger and older adults. Neurobiol Aging 48, 143–152. 10.1016/j.neurobiolaging.2016.08.007

De Cheveigné A, Arzounian D (2018) Robust detrending, rereferencing, outlier detection, and inpainting for multichannel data. NeuroImage 172, 903–912. 10.1016/j.neuroimage.2018.01.035

Dewar M, Alber J, Butler C, Cowan N, Della Sala S (2012) Brief Wakeful Resting Boosts New Memories Over the Long Term. Psychol Sci 23, 955–960. 10.1177/0956797612441220

Dewar M, Alber J, Cowan N, Della Sala S (2014) Boosting long-term memory via wakeful rest: intentional rehearsal is not necessary, consolidation is sufficient. PloS One 9, e109542. 10.1371/journal.pone.0109542

Dewar MT, Cowan N, Sala SD (2007) Forgetting Due to Retroactive Interference: A Fusion of Müller and Pilzecker’s (1900) Early Insights into Everyday Forgetting and Recent Research on Anterograde Amnesia. Cortex 43, 616–634. 10.1016/S0010-9452(08)70492-1

Evans FA, Stolwyk RJ, Wong D (2021) A Brief Period of Wakeful Rest after Learning Enhances Verbal Memory in Stroke Survivors. J Int Neuropsychol Soc 27, 929–938. 10.1017/S1355617720001307

Fatania J, Mercer T (2017) Nonspecific Retroactive Interference in Children and Adults. Adv Cogn Psychol 13, 314–322. 10.5709/acp-0231-6

Frossard J, Renaud O (2021) Permutation Tests for Regression, ANOVA, and Comparison of Signals: The permuco Package. J Stat Softw 99, 1–32. 10.18637/jss.v099.i15

Ghibaudo V, Turrel M, Granget J, Souilhol M, Garcia S, Plailly J, Buonviso N (2025) Pleasant odors specifically promote a soothing autonomic response and brain–body coupling through respiratory modulation. Sci Rep 15, 36417. 10.1038/s41598-025-20422-x

Gottselig JM, Hofer-Tinguely G, Borbély AA, Regel SJ, Landolt H-P, Rétey JV, Achermann P (2004) Sleep and rest facilitate auditory learning. Neuroscience 127, 557–561. 10.1016/j.neuroscience.2004.05.053

Gramfort A, Luessi M, Larson E, Engemann DA, Strohmeier D, Brodbeck C, Goj R, Jas M, Brooks T, Parkkonen L, Hämäläinen M (2013) MEG and EEG data analysis with MNE-Python. Front Neurosci 7, 267. 10.3389/fnins.2013.00267

Guinet V, Dantony E, Catenoix H, Nourredine M, Garcia S, Boulogne S, Leclercq M, Guen CL, Fourcaud-Trocmé N, Roy P, Ravel N, Rheims S (2026) Accelerated long-term forgetting in drug-resistant focal epilepsy: Insight from a prospective controlled study using a multimodal associative memory paradigm. medRxiv. 10.64898/2026.01.08.26343621

Humiston GB, Tucker MA, Summer T, Wamsley EJ (2019) Resting States and Memory Consolidation: A Preregistered Replication and Meta-Analysis. Sci Rep 9, 19345. 10.1038/s41598-019-56033-6

Humiston GB, Wamsley EJ (2018) A brief period of eyes-closed rest enhances motor skill consolidation. Neurobiol Learn Mem 155, 1–6. 10.1016/j.nlm.2018.06.002

Keeble L, Monaghan P, Robertson EM, Hannan S (2025) Slow-wave sleep as a key player in offline memory processing: insights from human EEG studies. Front Behav Neurosci 19, 1620544. 10.3389/fnbeh.2025.1620544

King O, Nicosia J (2022) The effects of wakeful rest on memory consolidation in an online memory study. Front Psychol 13, 932592. 10.3389/fpsyg.2022.932592

Leetham E, Watermeyer T, Craig M (2024) An online experiment that presents challenges for translating rest-related gains in visual detail memory from the laboratory to naturalistic settings. PLOS ONE 19, e0290811. 10.1371/journal.pone.0290811

Lüdecke D, Ben-Shachar M, Patil I, Waggoner P, Makowski D (2021) performance: An R Package for Assessment, Comparison and Testing of Statistical Models. JOSS 6, 3139. 10.21105/joss.03139

Maddirala AK, Veluvolu KC (2021) Eye-blink artifact removal from single channel EEG with k-means and SSA. Sci Rep 11, 11043. 10.1038/s41598-021-90437-7

Martini M, Martini C, Maran T, Sachse P (2018) Effects of post-encoding wakeful rest and study time on long-term memory performance. J Cogn Psychol 30, 558–569. 10.1080/20445911.2018.1506457

Martini M, Riedlsperger B, Maran T, Sachse P (2020) The Effect of Post-Learning Wakeful Rest on the Retention of Second Language Learning Material over the Long Term. Curr Psychol 39, 299–306. 10.1007/s12144-017-9760-z

Martini M, Sachse P (2020) Factors modulating the effects of waking rest on memory. Cogn Process 21, 149–153. 10.1007/s10339-019-00942-x

Martini M, Zamarian L, Sachse P, Martini C, Delazer M (2019) Wakeful resting and memory retention: a study with healthy older and younger adults. Cogn Process 20, 125–131. 10.1007/s10339-018-0891-4

MathWorks. (2024) MATLAB (R2024a) [Computer software]. The MathWorks, Inc. https://www.mathworks.com

Mednick SC, Cai DJ, Shuman T, Anagnostaras S, Wixted J (2011) An opportunistic theory of cellular and systems consolidation. Trends Neurosci 34, 504–514. 10.1016/j.tins.2011.06.003

Mercer T (2015) Wakeful rest alleviates interference-based forgetting. Memory 23, 127–137. 10.1080/09658211.2013.872279

Millar PR, Balota DA (2022) Wakeful Rest Benefits Recall, but Not Recognition, of Incidentally Encoded Memory Stimuli in Younger and Older Adults. Brain Sci 12, 1609. 10.3390/brainsci12121609

Posit team (2025) Rstudio: Integrated development environment for r. Posit Software, PBC. Boston, MA. http://www.posit.co/

Core Team (2025) R: A language and environment for statistical computing. R Foundation for Statistical Computing. Vienna, Austria. https://www.R-project.org/

Sacripante R, McIntosh RD, Della Sala S (2019) Benefit of wakeful resting on gist and peripheral memory retrieval in healthy younger and older adults. Neurosci Lett 705, 27–32. 10.1016/j.neulet.2019.04.026

Schlichting A, Bäuml K-HT (2017) Brief wakeful resting can eliminate directed forgetting. Mem 25, 254–260. 10.1080/09658211.2016.1153659

Schreiner T, Staudigl T (2020) Electrophysiological signatures of memory reactivation in humans. Philos Trans R Soc B Biol Sci 375, 20190293. 10.1098/rstb.2019.0293

Seban P, Šikl R, Prošek T, Urban K (2026) Wakeful rest and memory consolidation in an ecologically valid educational setting: no benefit over distractor tasks. Psychol Res 90, 42. 10.1007/s00426-026-02265-x

Sjøgård M, Baxter B, Mylonas D, Thompson M, Kwok K, Driscoll B, Tolosa A, Shi W, Stickgold R, Vangel M, Chu CJ, Manoach DS (2025) Hippocampal ripples predict motor learning during brief rest breaks in humans. Nat Commun 16, 1234. 10.1038/s41467-025-61136-y

Staresina BP, Bergmann TO, Bonnefond M, van der Meij R, Jensen O, Deuker L, Elger CE, Axmacher N, Fell J (2015) Hierarchical nesting of slow oscillations, spindles and ripples in the human hippocampus during sleep. Nat Neurosci 18, 1679–1686. 10.1038/nn.4119

Stawarczyk D, Majerus S, Van Der Linden M, D’Argembeau A (2012) Using the Daydreaming Frequency Scale to Investigate the Relationships between Mind-Wandering, Psychological Well-Being, and Present-Moment Awareness. Front Psychol 3, 363. 10.3389/fpsyg.2012.00363

Tucker MA, Humiston GB, Summer T, Wamsley E (2020) Comparing the Effects of Sleep and Rest on Memory Consolidation. Nat Sci Sleep 12, 79–91. 10.2147/NSS.S223917

Vallat R (2018) Pingouin: statistics in Python. J Open Source Softw 3, 1026. 10.21105/joss.01026

Vargha A, Delaney HD (2000) A Critique and Improvement of the CL Common Language Effect Size Statistics of McGraw and Wong. J Educ Behav Stat 25, 101–132. 10.3102/10769986025002101

Varma S, Daselaar SM, Kessels RPC, Takashima A (2018) Promotion and suppression of autobiographical thinking differentially affect episodic memory consolidation. PLOS ONE 13, e0201780. 10.1371/journal.pone.0201780

Varma S, Takashima A, Fu L, Kessels RPC (2019) Mindwandering propensity modulates episodic memory consolidation. Aging Clin Exp Res 31, 1601–1607. 10.1007/s40520-019-01251-1

Varma S, Takashima A, Krewinkel S, van Kooten M, Fu L, Medendorp WP, Kessels RPC, Daselaar SM (2017) Non-Interfering Effects of Active Post-Encoding Tasks on Episodic Memory Consolidation in Humans. Front Behav Neurosci 11, 54. 10.3389/fnbeh.2017.00054

Virtanen P et al. (2020) SciPy 1.0: fundamental algorithms for scientific computing in Python. Nat Methods 17, 261–272. 10.1038/s41592-019-0686-2

Walter M, Buonviso N, Plailly J (2024) Memory Consolidation During Wakeful Rest: Could Odors Play a Role?. 10.17605/OSF.IO/TVCX2

Walter M, Lacaze M, Garcia S, Buonviso N, Plailly J (2026) Supplements for ‘No Evidence for a Wakeful Rest Benefit on Associative Memory: A Within-Participant EEG Study’. 10.17605/OSF.IO/CJDE2

Wamsley EJ (2022) Offline memory consolidation during waking rest. Nat Rev Psychol 1, 441–453. 10.1038/s44159-022-00072-w

Wamsley EJ, Arora M, Gibson H, Powell P, Collins M (2023) Memory Consolidation during Ultra-short Offline States. J Cogn Neurosci 35, 1617–1634. 10.1162/jocn_a_02035

Wamsley EJ, Summer T (2020) Spontaneous Entry into an “Offline” State during Wakefulness: A Mechanism of Memory Consolidation? J Cogn Neurosci 32, 1714–1734. 10.1162/jocn_a_01587

Wang SY, Baker KC, Culbreth JL, Tracy O, Arora M, Liu T, Morris S, Collins MB, Wamsley EJ (2021) ‘Sleep-dependent’ memory consolidation? Brief periods of post-training rest and sleep provide an equivalent benefit for both declarative and procedural memory. Learn Mem 28, 195–203. 10.1101/lm.053330.120

Weng L, Yu J, Lv Z, Yang S, Jülich ST, Lei X (2025) Effects of wakeful rest on memory consolidation: A systematic review and meta-analysis. Psychon Bull Rev 32, 1937–1968. 10.3758/s13423-025-02665-x

Zhang H, Fell J, Axmacher N (2018) Electrophysiological mechanisms of human memory consolidation. Nat Commun 9, 4103. 10.1038/s41467-018-06553-y

