## Supplementary File for "No Evidence for a Wakeful Rest Benefit on Associative Memory: A Within-Participant EEG Study"

#### 1 Original Study Protocol

The original study was designed to study the impact of diffusing a personally pleasant essential oil during the two post-learning conditions: Rest and Distraction task. For this experiment, 60 participants were recruited and divided into two groups: O+ (i.e., with an odor diffused in the room during the post-learning conditions) and O- (i.e., without odor). Only the specificities related to the O+ group or the steps not described in the article are presented here.

*Session 1*

The aim of the first session was to select the favorite odor for each participant of the O+ group among 10 essential oils. The first session took place as follows:

For the O+ group:

*European Test of Olfactory Capacities* (ETOC, Thomas-Danguin et al., 2003): An olfactory test designed to detect olfactory deficits, combining a supraliminal detection task and an identification task. Only normosmic participants were included in this study.

*Daydreaming Frequency Scale* (DDFS, Stawarczyk et al., 2012).

*Odor Awareness Scale* (OAS, Smeets et al., 2008): This scale assesses individual differences in the importance and awareness of positive and negative odors in the environment.

*Selection of the favorite odor:*

Step 1: Odors were presented to participants on smelling strips in a randomized order. For each odor, participants decided whether it was either pleasant or unpleasant (hedonicity) and relaxing or stimulating (relaxation).

Step 2: For each odor categorized as both pleasant and relaxing, participants evaluated the following odor features on continuous scales:

- *Hedonicity*: "Indicate how much you like this smell" (from -5: very unpleasant to 5: very pleasant, with 0: neutral)

- *Relaxation*: "Indicate how much this smell relaxes you" (from -5: Energizing / Revitalizing / Dynamizing to 5: Relaxing / Soothing / Calming with 0: no effect)

- *Attraction*: "Indicate how much you would like to spend 10 minutes in a room with this smell" (from 0: not at all to 10: extremely)

Participants were free to explore the odors for as long as needed. If a participant did not classify any odor as both pleasant and relaxing, they were not selected for Session 2. If only one odor was classified as both pleasant and relaxing, that odor was selected for Session 2. If more than one odor was classified as both pleasant and relaxing, we selected the favorite odor based on the highest attraction score and confirmed it with the hedonicity and relaxation scores.

For the O- group:

Because we did not want the participants in the O- group to expect any odor presentation in the experiment, Session 1 slightly differed. First, their olfactory abilities were not assessed using the ETOC, and they did not complete the OAS but only the DDFS. Second, instead of the odor selection step, participants evaluated images of the 10 plants used in our selected essential oils. To do so, pictures of plants were randomly displayed on a computer screen and evaluated by participants with the same two steps as the O+ group. This evaluation of plant images enabled us to approximate the O+ condition without informing this group about odors.

*Session 2*

The second session took place a maximum of one week after the first.

*Sleep Questionnaires*

First, because of the well-known negative impact of sleepiness on memory performance, and to test a potential modulatory effect, participants indicated their level of sleepiness by answering three questions:

- “How many hours did you sleep last night?”

- “When you woke up this morning, did you feel like you had slept well?” (choice between: “yes”, “rather yes”, “rather no”, “no”)

- Karolinska Sleepiness Scale (Åkerstedt and Gillberg, 1990): The scale evaluates the subjective level of sleepiness at a particular time of day. Participants are asked to assess which level best reflects their psychophysical state over the last 10 minutes from 0 (extremely awake) to 9 (very sleepy, a lot of effort to stay awake, fights sleep).

*Setup and basal recording of electrophysiological activity*

We recorded breathing, cardiac and brain electrical activity for five minutes with participants’ eyes open.

*Memory Task Training*

Participants first underwent training with a subset of three trials to ensure task understanding.

*Relaxation Assessment*

Subjective relaxation was assessed using the short-STAI (Marteau and Bekker, 1992) along with two custom questions:

- How relaxed do you feel right now (physically, mentally)? (from -5: anxious, stressed, tensed to 5: relaxed, appeased, calmed with 0: neutral)

- How awake do you feel right now (mentally, physically) ? (from -5: tired, drowsy, sleepy to 5: energized, alert, excited, with 0: neutral)

*Associative Memory Task*

The memory task is described in the method section (see Sections 2.2 and 2.3).

*Post-Learning Sessions*

For the O+ group, the selected pleasant odor was diffused in the room during Rest and Distraction task, using a nebulization diffuser (Litsea model, Compagnie des Sens, Lyon, France).

*Final Questionnaires*

At the end of the memory task and after the equipment was removed, participants completed a questionnaire regarding their mental activities during the two post-learning periods: Rest and Distraction (see 2.3.1. Questionnaires). Finally, participants answered a last questionnaire with the following questions:

- "How difficult did you find the task of musical comparison?" (from 0: very easy to 10: very difficult)

- “How relaxing did you find the odor?” (-5: Energizing, Revitalizing, Dynamizing; 0: no effect; 5: Relaxing, Soothing, Calming)

- “How did you like the smell?” (-5: very unpleasant; 0: neutral; 5: very pleasant)

- “How much of a sensation of temperature did this smell generate?” (-5: cold; 0: neutral, 5: warm)

- “How much of a sensation of irritation did this smell generate?” (0: not at all; 10: extremely)

- “During the moment of rest, about the emotions you felt: how intense were these emotions?” (0: not very intense; 10: very intense)

- “During the moment of rest, regarding the memories recalled: how intense were these memories?” (0: not very intense; 10: very intense)

- “Please list the words that the smell evoked in you during the experiment.”

- "If you've identified the odor, please write down its name. Then, briefly explain the reasons why you preferred this smell (if any)."

- “During the moment of rest, how focused was your attention on your breathing?” (0: not at all; 10: extremely)

At the end of the experiment, all the item pictures were displayed one by one, and participants were asked to name them aloud as they did during the recall phases of the memory task. This step was done to ensure consistency with the experimenter's naming and to help prevent scoring errors. Before leaving, participants were asked if they understood the objectives and hypotheses of the study.

### 2 Analyses – supplementary informations

##### Implementation of Generalized Linear Mixed Models – model selection

Models were fitted using the *glmmTMB* package (version 1.1.13; Brooks et al., 2017) in R. Two random-effects structures were compared for each outcome: a random intercept per participant only and a random intercept with a by-participant random slope for the condition. Models performance were evaluated and compared using the *performance* package (version 0.15.2; Lüdecke et al., 2021). Model selection proceeded in two steps. A likelihood ratio test (LRT, using an ANOVA) assessed whether the added complexity of the slope term was statistically justified. If the LRT was non-significant, the more parsimonious random-intercept model was retained.. Model singularity and convergence were assessed using *check_convergence()* and *check_singularity()* functions, respectively. Model validity was further verified using the *check_predictions()* function. Regularizing priors were used to aid convergence if needed. Once all criteria were satisfied, estimated marginal means were computed and contrasts of interest were examined using the *emmeans* package (version 1.11.2.8; Lenth and Piaskowski, 2025).

##### Comparison of Analyses Pipelines – methodological concerns

To assess whether the choice of memory score influenced our results, we swapped the memory scores between pipelines: Relative Memory Change was applied within the replication-inspired pipeline, and Global Memory Change within the data-specific pipeline. Both scores yielded comparable correlations (e.g., Supplementary Figure 8), suggesting that the observed differences are more likely attributable to EEG preprocessing and processing decisions than to the choice of memory score.

Among the preprocessing parameters examined, window duration for PSD computation appears to play a meaningful role, particularly for slow oscillation power estimation. We considered 4-second Hanning windows, as used in Brokaw et al. (2016), insufficiently long to reliably capture SO power. Our primary motivation for the data-specific analysis was therefore to re-examine the correlation between SO power during Rest and memory performance using 20-second windows. Applying 20-second windows within the replication-inspired pipeline did not alter the direction or significance of the results, yet produced non-negligible changes in SO power estimates (Supplementary Figure 9), confirming that window length is a relevant parameter for this frequency band.

The choice of offline re-referencing scheme also proved to be a key factor. This has previously been shown to substantially affect EEG analyses (Hu et al., 2018), and our data confirm this sensitivity in some cases: switching the re-referencing method between average and mastoid-like (TP9/TP10) references nearly reversed the direction of certain correlations, without however reaching significance in either case (Supplementary Figure 10). This finding underscores the importance of re-referencing choices in EEG research, particularly when comparing results across studies that differ on this parameter.

Other preprocessing decisions not systematically tested here may further modulate results, including the maximum frequency cutoff used for PSD computation and the normalization of PSD by the aperiodic 1/f component (Donoghue et al., 2020) among others.

### 3 Supplementary Results

<u>Distraction task</u>

Distraction task difficulty was rated as 7.25 (± 1.74 MAD, min: 2.88, max: 10). Performances were of 39 (up to 54) in median (± 4.45 MAD, min: 26, max: 50), with a median response time of

4.45 (± 0.4 MAD, min: 4.05, max: 8.08) s. No correlation was observed between *Global Memory Change nor Relative Memory Change* in the Distraction condition and Perceived task difficulty, performances, nor response time (Supplementary Figure 7).

### 4 Supplementary Figures

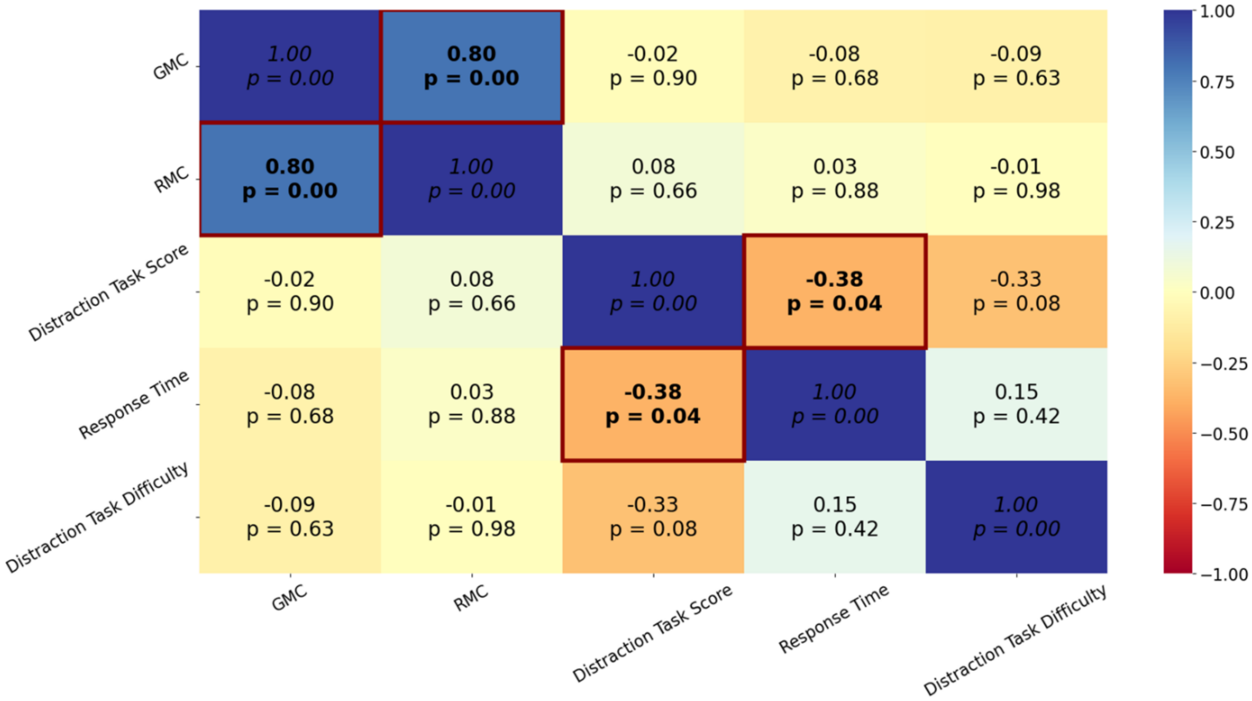


**Supplementary Figure 1 – Correlation matrix of distraction task measures with memory performance.** Spearman *r* coefficient and corresponding *p*-values are indicated for each pair of memory performance metrics (*Global Memory Change (GMC)* and *Relative Memory Change (RMC))* and distraction task measures (score, response time, and perceived difficulty). Matrix cells are color-coded from dark blue (*r* = 1, strong positive relationship) to red (*r* = -1, strong negative relationship). Cells indicating significant correlations are framed in dark red with the text in bold.

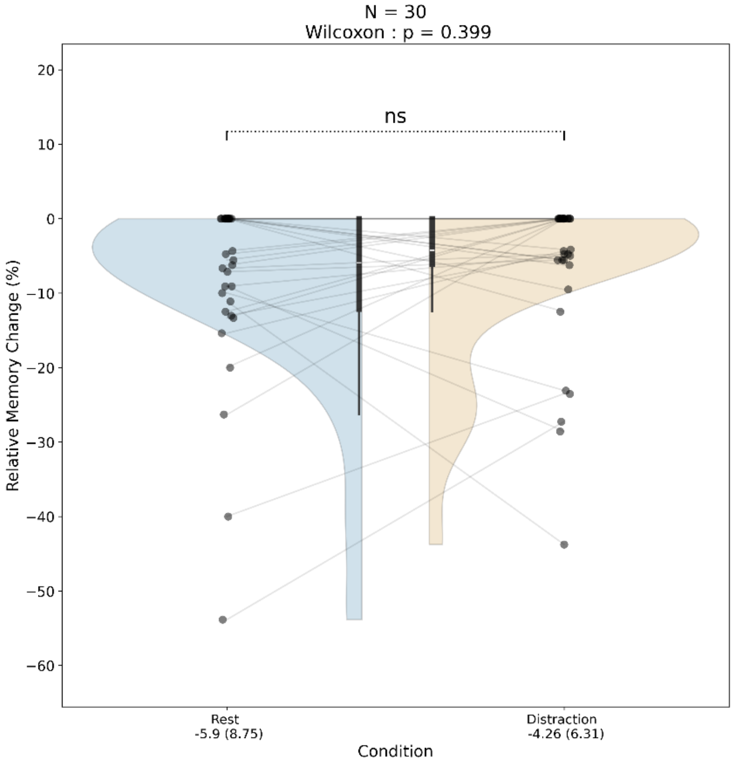


**Supplementary Figure 2 – Effects of experimental conditions on memory performance (*Relative Memory Change*).** Violin plots show the full data distribution (ranging from minimum to maximum) of *Relative Memory Change* for each condition with individual data points and lines connecting those from the same participant. Inner boxplots indicate central tendancies (whiskers = 1.5 times the interquartile range beyond the box; box = first quartile, median, third quartile); ns: non-significant.

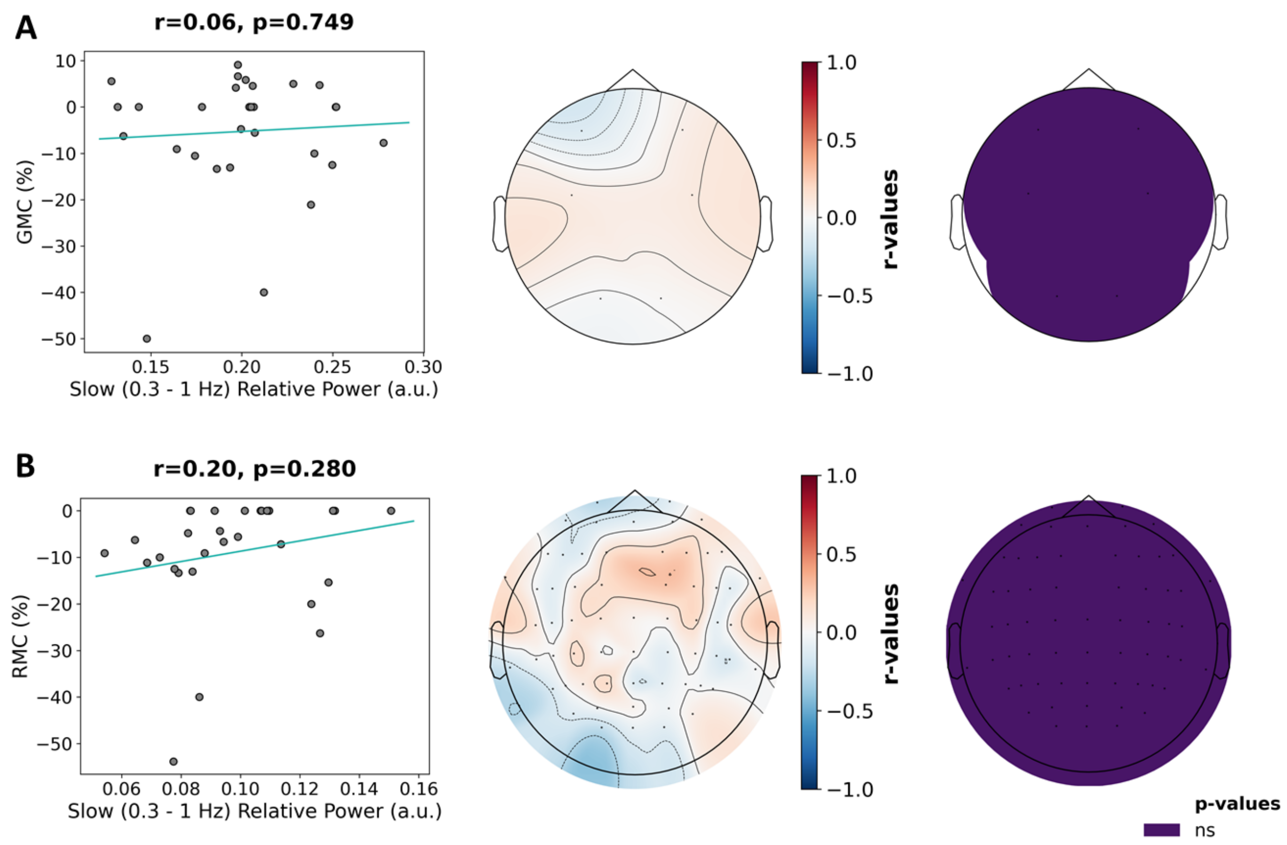


**Supplementary Figure 3 – Correlation between slow oscillation power and memory performance for the Rest condition.** (A) Replication-inspired analysis restricted to the six channels used in Brokaw et al. (2016) with 4-second windows for PSD computation, using TP9 and TP10 channels as references and Global Memory Change as memory performance (B) PSD analysis conducted across 62 channels with 20-second Hanning windows using an average reference and Relative Memory Change as memory performance. For both analyses: Left panel: linear regression on original non-ranked values of memory performance against slow oscillations (SO, 0.3 – 1 Hz) power. Middle panel: topomap of Spearman correlation coefficients (r-values) per channel. Right panel: topomap of channels with significant correlations with memory performance. No significant correlation was observed between SO power and memory performance in either analysis.

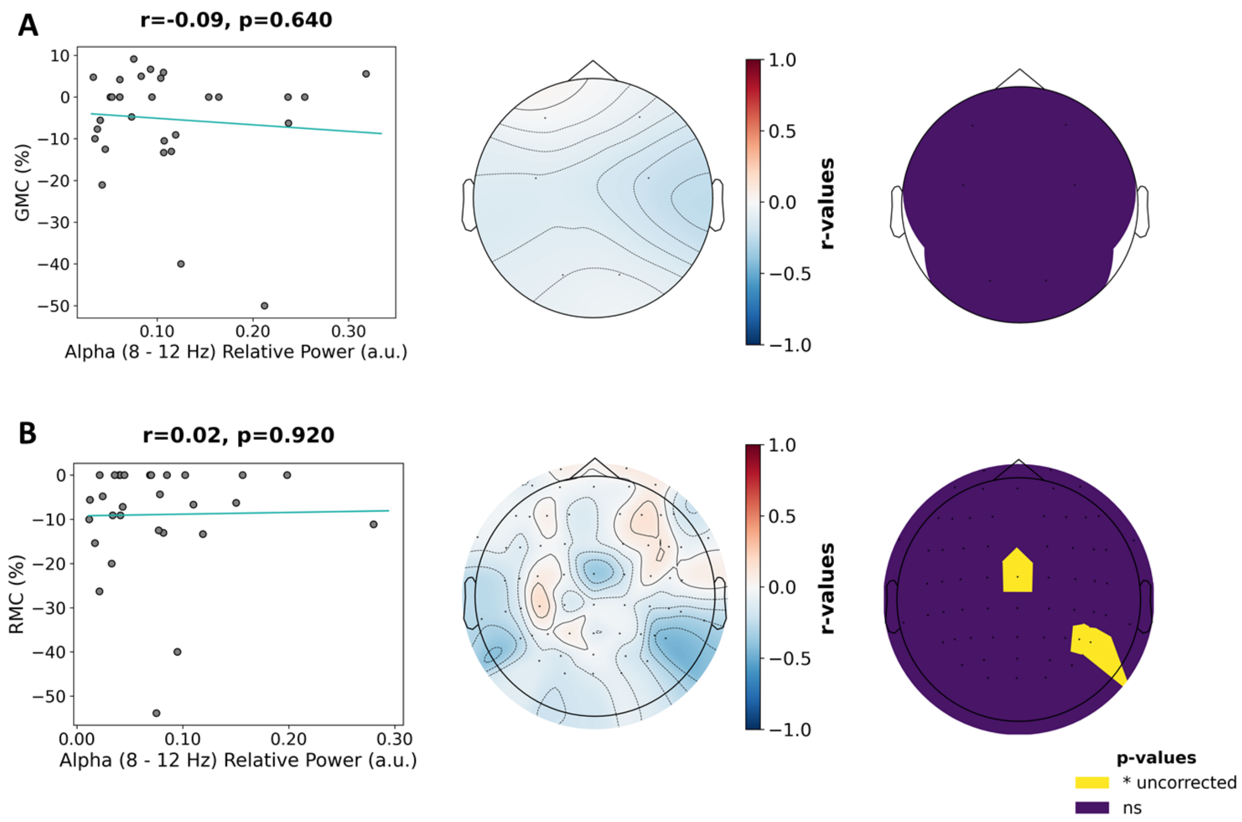


**Supplementary Figure 4 – Correlation between alpha power and memory performance for the Rest condition.** (A) Replication-inspired analysis restricted to the six channels used in Brokaw et al. (2016) with 4-second windows for PSD computation, using TP9 and TP10 channels as references and Global Memory Change as memory performance (B) PSD analysis conducted across 62 channels with 20-second Hanning windows using an average reference and Relative Memory Change as memory performance. For both analyses: Left panel: linear regression original non ranked of memory performances against alpha (8 – 12 Hz) power. Middle panel: topomap of Spearman correlation coefficients (r-values) per channel. Right panel: topomap of channels with significant correlations with memory performance. Channels are color-coded as follows: indigo, non-significant; yellow, significant uncorrected. No significant correlation was observed between alpha power and memory performance in either analysis.

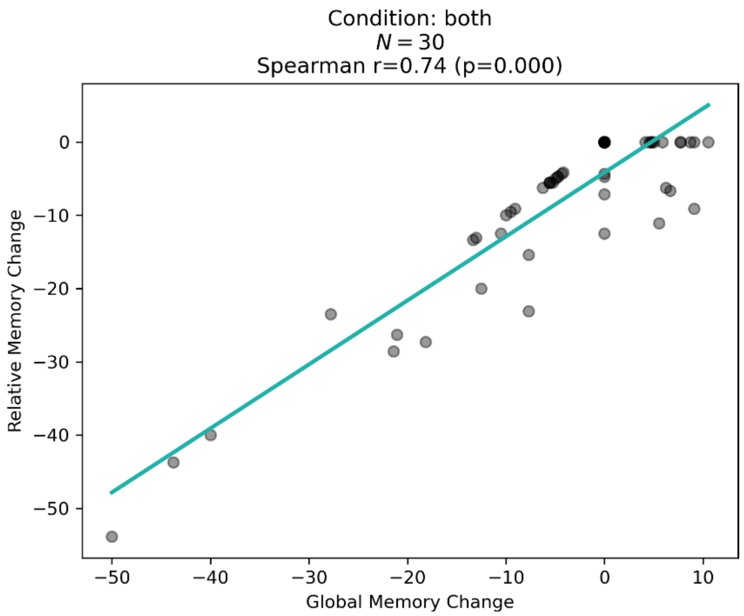


**Supplementary Figure 5 – Correlation between *Global Memory Change* and *Relative Memory Change.*** The blue line represents the regression fit.

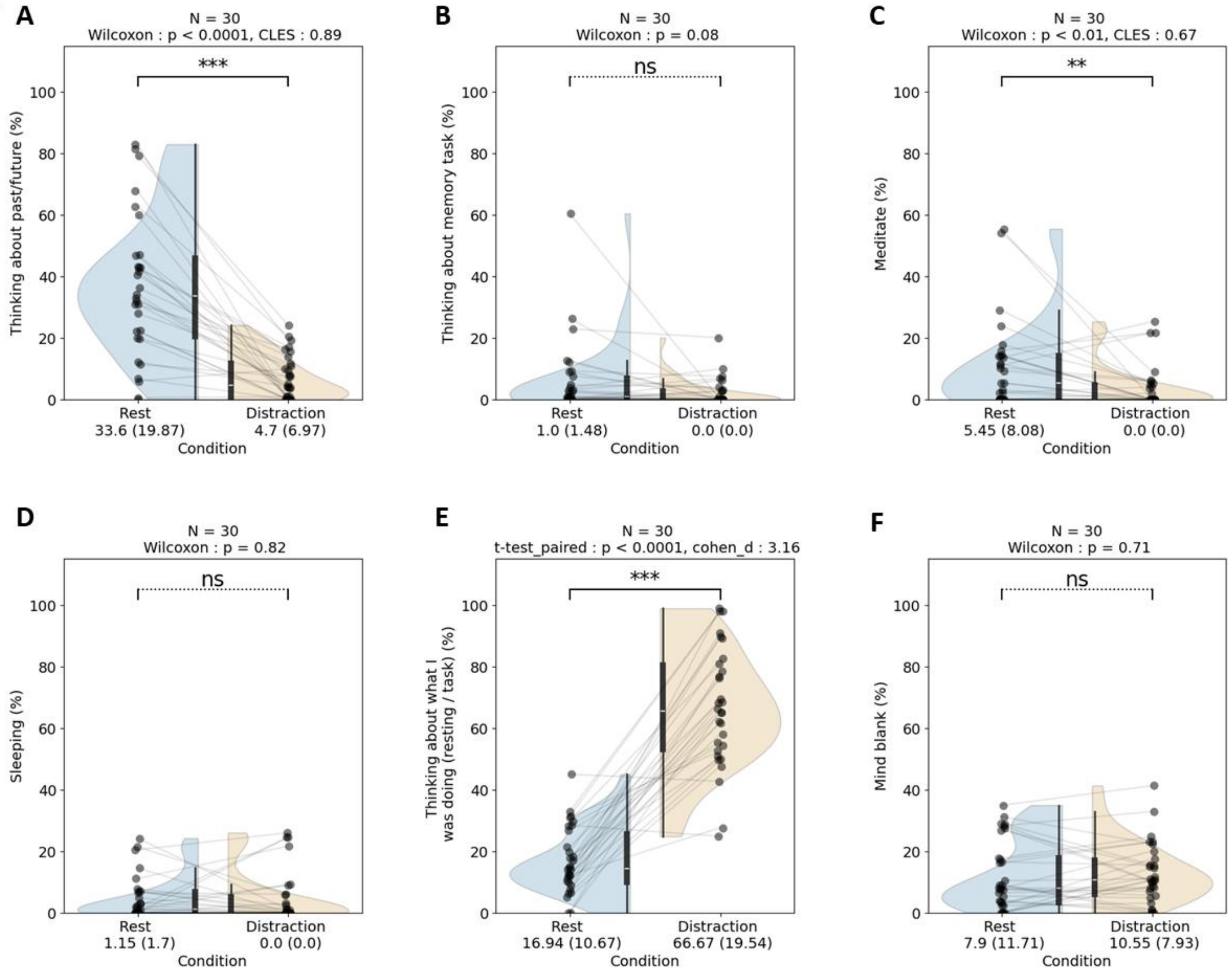


**Supplementary figure 6 – Mental activities during post-learning conditions.** Each subplot corresponds to the comparison of the percentage of time engaged in one of the proposed mental activities between Rest and Distraction: (A) Thinking about the past/future, (B) Thinking about the memory task, (C) Meditating, (D) Sleeping, (E) Thinking about what I was doing, (F) Having a mind blank. Violin plots show the full data distribution (ranging from minimum to maximum). Individual data points are superimposed on the plots, with lines connecting those from the same participant. Inner boxplots indicate central tendancies (whiskers = 1.5 times the interquartile range beyond the box; box = first quartile, median, third quartile). CLES = Common Language Effect Size; *** = $p < 0.001$ ; ** = $p < 0.01$; ns = non-significant.

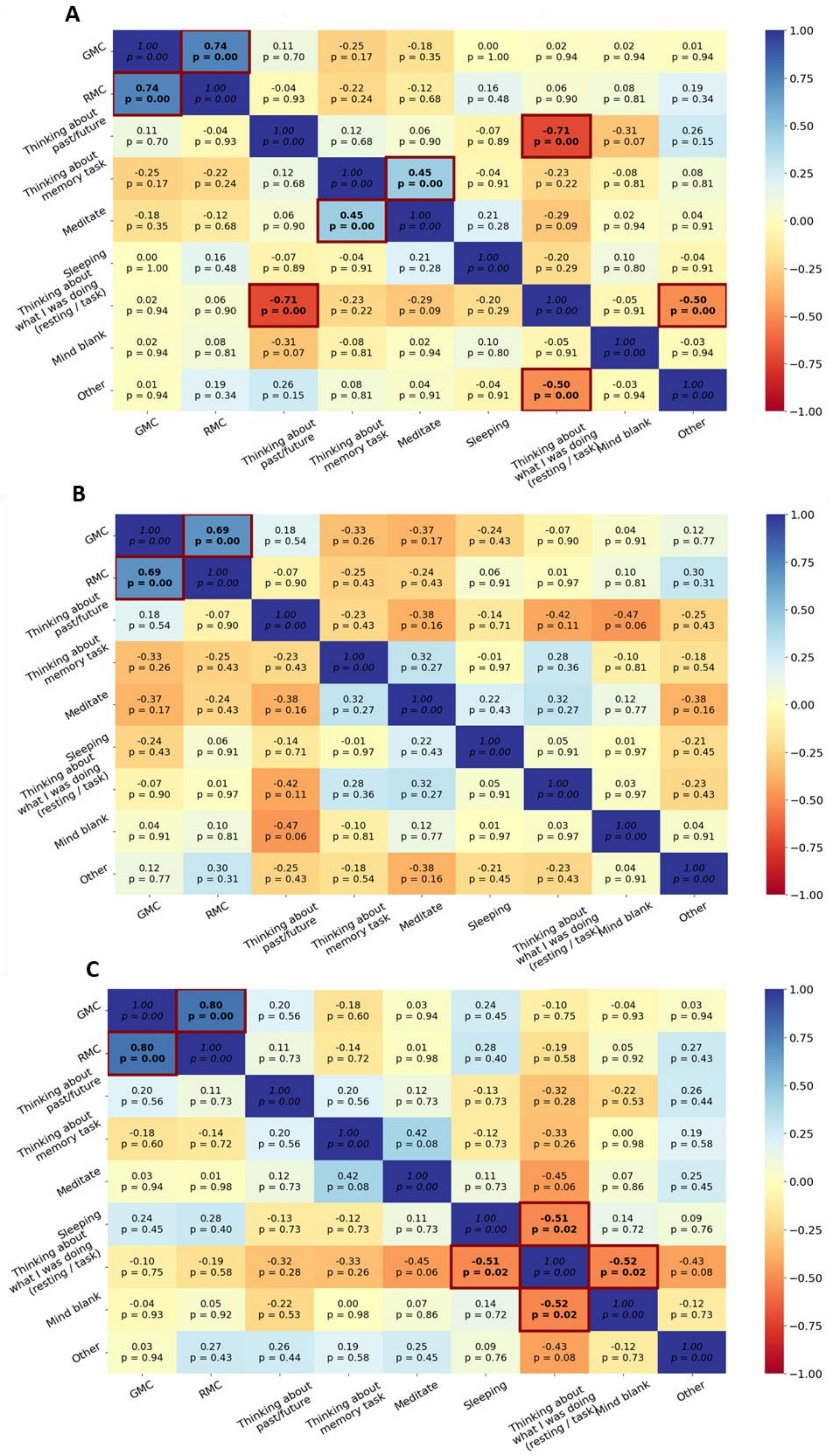
A
B
C
GMC
RMC
Thinking about past/future
Thinking about memory task
Meditate
Sleeping
Thinking about what I was doing (resting / task)
Mind blank
Other
1.00
0.75
0.50
0.25
0.00
−0.25
−0.50
−0.75
−1.00

**Supplementary Figure 7 – Correlation matrices between memory performance and mental activities during post-learning conditions.** Spearman *r* coefficient and corresponding *p*-values (corrected for multiple comparisons using FDR) are indicated for each pair of memory performance metrics (*Global Memory Change* and *Relative Memory Change)* and each mental activity (Thinking about past/future, Thinking about memory task, Meditate, Sleeping, Thinking about what I was doing (resting/task), Mind blank, Other). Matrix cells are color coded from dark blue (r = 1, strong positive relationship) to red (r = -1, strong negative relationship). Cells indicating significant correlations are framed in dark red with the text in bold. (A) Both conditions together. (B) Only Rest condition. (C) Only Distraction condition.

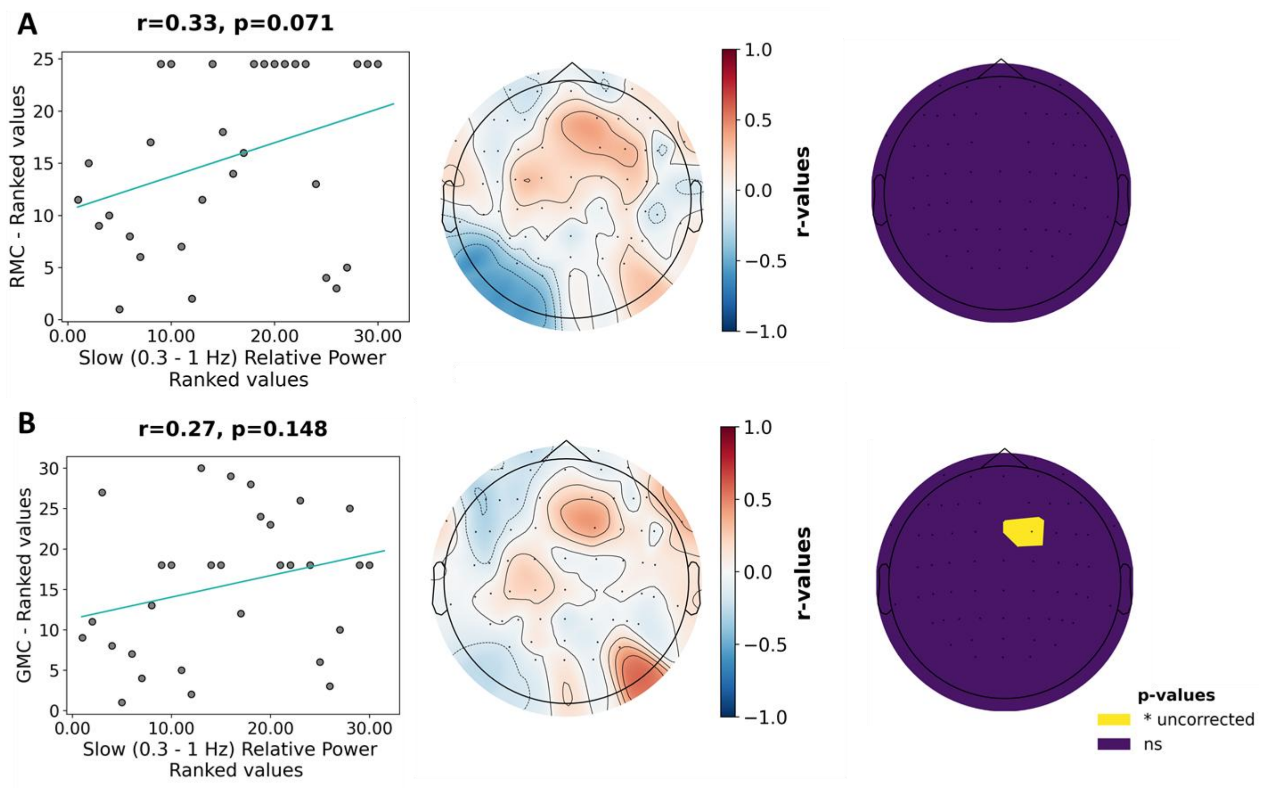


**Supplementary Figure 8 – Comparison of correlations between slow oscillation power and memory performance when memory scores are reversed.** PSD analysis conducted across 62 channels with 20-second Hanning windows using an average reference and Global Memory Change as memory performances. For both analyses: Left panel: linear regression on ranked values of memory performance against slow oscillations (SO, 0.3 – 1 Hz) power. Middle panel: topomap of Spearman correlation coefficients (r-values) per channel. Right panel: topomap of channels with significant correlations with memory performances. (A) using Relative Memory Change as memory performances. (B) using Global Memory Change as memory performances. Correlations look very similar even when reversing the memory performance score used in replication-inspired and data-specific analyses.

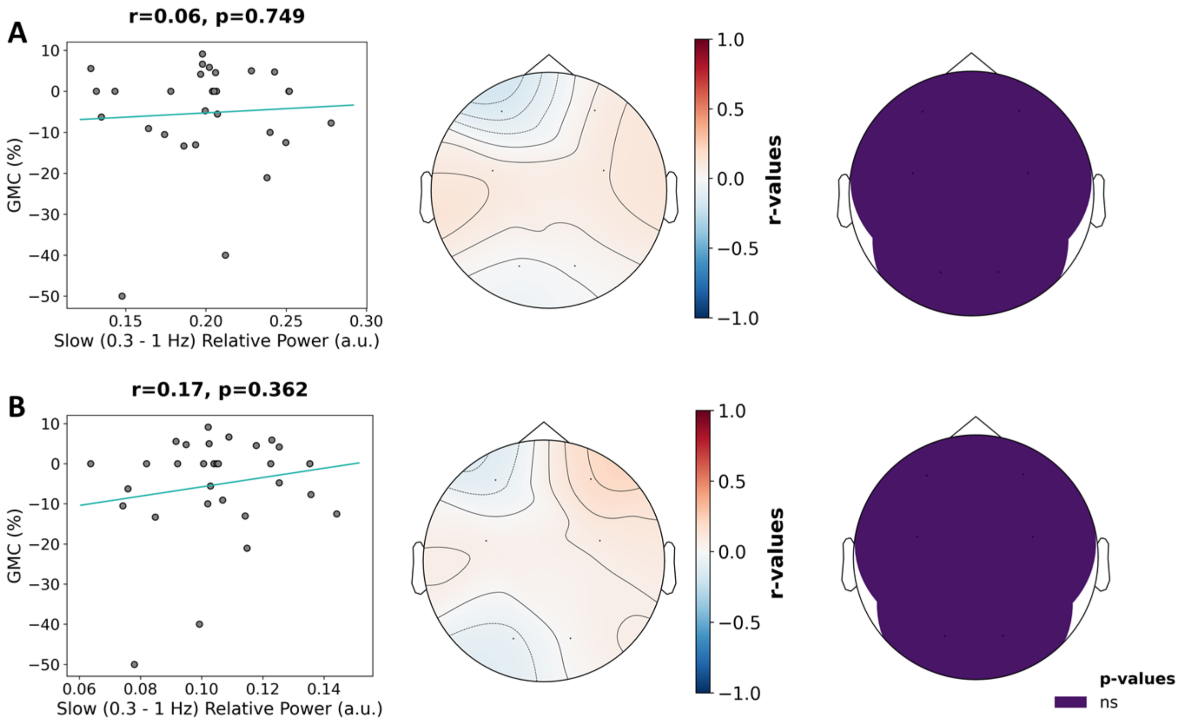


**Supplementary Figure 9 – Comparison of correlations between slow oscillation power and memory performance when window durations for PSD computation are reversed.** For both analyses: Left panel: linear regression on original non-ranked values of memory performance against slow oscillations (SO, 0.3 – 1 Hz) power. Middle panel: topomap of Spearman correlation coefficients (r-values) per channel. Right panel: topomap of channels with significant correlations with memory performances. (A) PSD analysis conducted across 6 channels with 4-second Hanning windows using TP9 and TP10 channels as references and Global Memory Change as memory performances. (B) PSD analysis conducted across 6 channels with 20-second Hanning windows using TP9 and TP10 channels as references and Global Memory Change as memory performances. While not changing the significance and direction of the relationship, window size for PSD computation seems to play a non-negligible role, especially for SO power PSD computation.

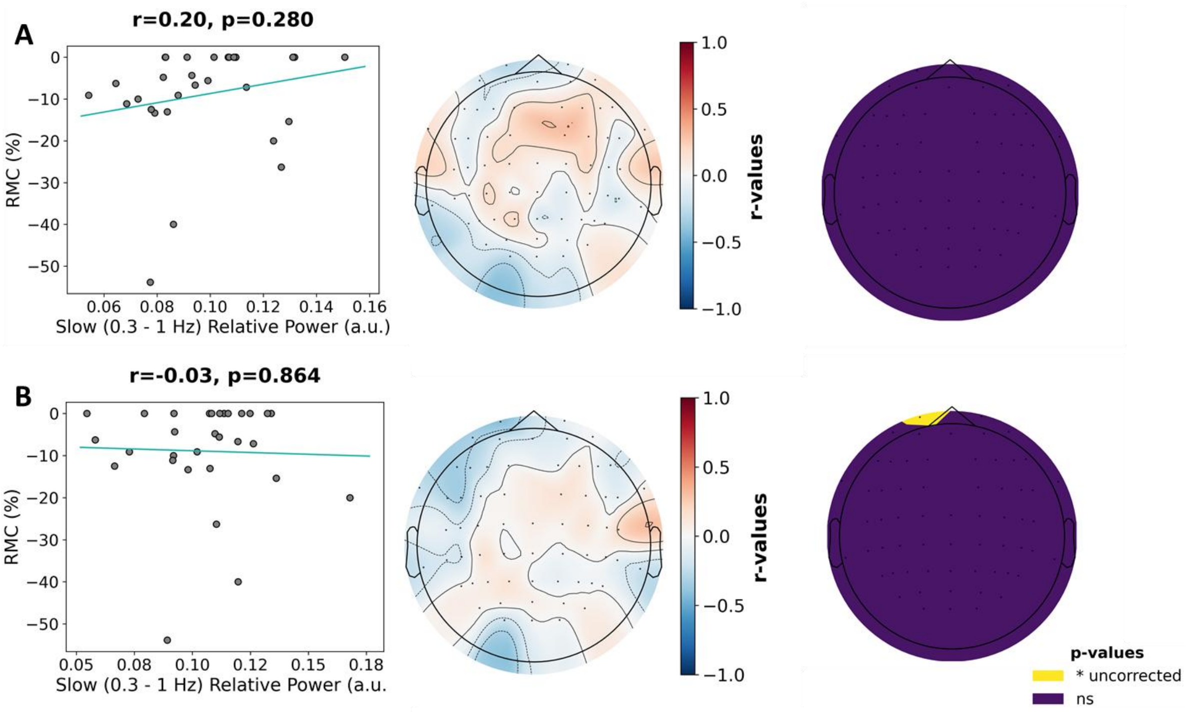


**Supplementary Figure 10 – Comparison of correlations between slow oscillation power and memory performance when reversing the re-referencing method.** For both analyses: Left panel: linear regression on original non-ranked values of memory performance against slow oscillations (SO, 0.3 – 1 Hz) power. Middle panel: topomap of Spearman correlation coefficients (r-values) per channel. Right panel: topomap of channels with significant correlations with memory performances. (A) PSD analysis conducted across 62 channels with 20-second Hanning windows, an average reference and Relative Memory Change as memory performances. (B) PSD analysis conducted across 62 channels with 20-second Hanning windows using TP9 and TP10 channels as references and Relative Memory Change as memory performances. The direction of correlations is almost inverted when the referencing method is changed, although none are significant.

#### 5 References


Åkerstedt T, Gillberg M (1990) Subjective and Objective Sleepiness in the Active Individual. Int J Neurosci 52:29–37.

Brokaw K, Tishler W, Manceor S, Hamilton K, Gaulden A, Parr E, Wamsley EJ (2016) Resting state EEG correlates of memory consolidation. Neurobiol Learn Mem 130:17–25.

Brooks ME, Kristensen K, Benthem KJ van, Magnusson A, Berg CW, Nielsen A, Skaug HJ, Mächler M, Bolker BM (2017) glmmTMB Balances Speed and Flexibility Among Packages for Zero-inflated Generalized Linear Mixed Modeling. R J 9:378–400.

Donoghue T, Haller M, Peterson EJ, Varma P, Sebastian P, Gao R, Noto T, Lara AH, Wallis JD, Knight RT, Shestyuk A, Voytek B (2020) Parameterizing neural power spectra into periodic and aperiodic components. Nat Neurosci 23:1655–1665.

Hu S, Lai Y, Valdes-Sosa PA, Bringas-Vega ML, Yao D (2018) How do reference montage and electrodes setup affect the measured scalp EEG potentials? J Neural Eng 15:026013.

Lüdecke D, Ben-Shachar M, Patil I, Waggoner P, Makowski D (2021) performance: An R Package for Assessment, Comparison and Testing of Statistical Models. J Open Source Softw 6:3139.

Marteau TM, Bekker H (1992) The development of a six-item short-form of the state scale of the Spielberger State—Trait Anxiety Inventory (STAI). Br J Clin Psychol 31:301–306.

Smeets MAM, Schifferstein HNJ, Boelema SR, Lensvelt-Mulders G (2008) The Odor Awareness Scale: A New Scale for Measuring Positive and Negative Odor Awareness. Chem Senses 33:725–734.

Stawarczyk D, Majerus S, Van Der Linden M, D'Argembeau A (2012) Using the Daydreaming Frequency Scale to Investigate the Relationships between Mind-Wandering, Psychological Well-Being, and Present-Moment Awareness. Front Psychol 3, 363.

Thomas-Danguin T, Rouby C, Sicard G, Vigouroux M, Farget V, Johanson A, Bengtzon A, Hall G, Ormel W, De Graaf C, Rousseau F, Dumont J-P (2003) Development of the ETOC: a European test of olfactory capabilities. Rhinology 41:142–151.
